# Polyphosphate mobilization influences the ability of *Cryptococcus neoformans* to cause disease in a murine model of cryptococcosis

**DOI:** 10.1101/2024.05.28.596173

**Authors:** Kabir Bhalla, Eddy Sánchez León-Hing, Yu-Hsuan Huang, Victoria French, Guanggan Hu, Jennifer Wang, Matthias Kretschmer, Xianya Qu, Raphaell Moreira, E. Johan Foster, Pauline Johnson, James W. Kronstad

**Affiliations:** Department of Microbiology and Immunology, Michael Smith Laboratories, University of British Columbia, Vancouver BC, Canada; Department of Microbiology and Immunology, Life Sciences Institute, University of British Columbia, - Vancouver BC, Canada; Department of Chemical and Biological Engineering, BioProducts Institute, University of British Columbia, Vancouver BC, Canada

## Abstract

*Cryptococcus neoformans*, an invasive basidiomycete fungal pathogen, causes one of the most prevalent, life-threatening diseases in immunocompromised individuals and accounts for ∼15% of AIDS-associated deaths. A dire need for the development of novel antifungal drugs, vaccines, and improved diagnostics has emerged with the increased frequency of fungal infections. Therefore, understanding the pathogenesis of *C. neoformans* and its interactions with the host immune system is critical for the development of therapeutics against cryptococcosis. Previous research demonstrated that *C. neoformans* cells lacking polyphosphate (polyP), an immunomodulatory polyanionic storage molecule, display altered cell surface architecture. However, the relevance of surface changes and the role of polyP in the virulence of *C. neoformans* remain unclear. Here we show that mutants lacking the polyphosphatases (Xpp1 and Epp1) are attenuated for virulence in a murine inhalational model of cryptococcosis, demonstrate reduced proliferation in host tissue, and provoke an altered immune response. An analysis of mutants lacking the polyphosphatases and the Vtc4 protein for polyP synthesis indicated that the Xpp1 and Epp1 contribute to the organization of the cell surface, virulence factor production, the response to stress, and mitochondrial function. Overall, we conclude that polyP mobilization plays a multifaceted role in the pathogenesis of *C. neoformans*.

**Author Summary:** *Cryptococcus neoformans* causes one of the most prevalent fungal diseases in people with compromised immune systems and accounts for 15-20% of AIDS-associated deaths worldwide. The continual increase in the incidence of fungal infections and limited treatment options necessitate the development of new antifungal drugs and improved diagnostics. Polyphosphate (polyP), an under-explored biopolymer, functions as a storage molecule, modulates the host immune response, and contributes to the ability of many fungal and bacterial pathogens to cause disease. However, the role of polyP in cryptococcal disease remains unclear. In this study, we report that the enzymes that regulate polyP synthesis and turnover contribute to the virulence of *C. neoformans* in a mouse model of cryptococcosis. The polyphosphatases, Xpp1 and Epp1, influenced the survival of *C. neoformans* in macrophages and altered the host immune response. The loss of Xpp1 and Epp1 led to changes in cell surface architecture, cell size, impaired growth, and defects in both mitochondrial function and the stress response of *C. neoformans.* Thus, our work establishes polyP as a key factor in the disease caused by *C. neoformans*, and identifies polyP mobilization as a novel target to support new therapeutic approaches.

## Introduction

Fungi are underappreciated and neglected as major pathogens of humans despite over a billion individuals impacted each year worldwide (1–3). As an example, cryptococcal disease is responsible for ∼300,000 cases of meningoencephalitis each year and is the second leading cause of AIDS-associated deaths in sub-Saharan Africa (2,4). Acquired by inhalation of basidiospores or desiccated yeast cells, *C. neoformans* has a propensity to disseminate to the central nervous system and cause life-threatening disease in immunocompromised individuals (5). Currently, the treatment for cryptococcosis involves long term therapy, but limited antifungal agents are available (6). The continual increase in the incidence of fungal infections and the notoriously difficult nature of treatment due to host toxicity and anti-cryptococcal drug resistance highlights the critical need to develop new antifungal drugs and improved diagnostics (2). Cryptococcal disease is also a more general concern for public health because *Cryptococcus gattii*, a closely related sibling species with shared virulence factors, can cause cryptococcosis in immunocompetent individuals without any underlying health conditions (7,8).

As a facultative intracellular pathogen, *C. neoformans* employs several host damage mechanisms and virulence factors that help it evade host immunity, persist in the intracellular environment, and resist the unfavourable conditions of the phagolysosome (8,9). One of the major virulence factors is the polysaccharide capsule which modulates immune responses and prevents recognition and phagocytosis by masking immunoreactive pathogen associated molecular patterns (PAMPs) such as chitin and chitosan (8,10,11). Glucuronoxylomannan (GXM) and glucuronoxylomannogalactan (GXMGal), the two major components of the capsule, have additional functions in depletion of complement and inhibition of neutrophil migration (12). Moreover, the capsule is involved in protecting the fungus from oxidative stresses (11). In addition, *C. neoformans* produces melanin, an immunogenic natural pigment and antioxidant, which provides further protection against oxidative damage and fluctuations in temperature (11). Production of melanin also confers resistance to antifungal drugs such as amphotericin B and contributes to fungal dissemination to the brain tissue (13,14). Lastly, titanization, or formation of morphologically giant cells characterized by thicker cell walls and large vacuoles, is a unique virulence factor which allows *C. neoformans* to evade phagocytosis (15,16).

Upon inhalation, *C. neoformans* cells colonize the lung tissue and encounter resident immune cells (17,18). β-glucans and chitin, as well as chitosan present on the cell wall act as fungal PAMPs and are recognized by immune cells through pattern recognition receptors (PRRs) such as Toll-like receptors (TLRs) and C-type lectin receptors (CLRs) to trigger appropriate cytokine and phagocytic responses (10,19,20). Alveolar macrophages, the first line of defense, detect and phagocytose cryptococcal cells, and recruit monocytes and dendritic cells (DCs) to the lung (18,21). These recruited DCs also internalize yeast cells, and present antigens to naïve CD4+ T cells in the lymph nodes (18,21). As with other intracellular pathogens, the polarization and induction of IFNγ-, IL-6-, and IL-17-secreting T helper 1 (Th1) and T helper 17 (Th17) cells promote pathogen clearance (22). In fact, mice lacking IFNγ receptors display high susceptibility to cryptococcosis and have increased fungal burdens (23). Furthermore, mice deficient in both Th1 and Th17 responses display heightened mortality to cryptococcal infections compared with mice deficient in only Th1 responses (24). The commonly studied wild-type *C. neoformans* strain, H99, is known to induce a non-protective Th2 immune response through production of IL-4 in BALB/c mice (8). Th2 responses are maladaptive and are commonly associated with promoting fungal dissemination and proliferation (25).

Polyphosphate (polyP), an under-explored, polyanionic inorganic biopolymer, serves as a phosphorus and energy storage molecule (26). In fungi, polyP is found in different cellular compartments where it complexes with various inorganic and organic cations (27). Recent findings also identified involvement of this polyanion in osmoregulation, stress adaptation, cell membrane formation, biofilm development, quorum sensing, and protein targeting (28–31). PolyP is also known to contribute to virulence in many bacterial pathogens such as *Salmonella* and *Mycobacterium* (27). More specifically, prokaryotic polyphosphates inhibit the polarization of M1 macrophages, suppress inducible nitric oxide synthase (iNOS), and impede major histocompatibility complex class II (MHC II) activation in the host (28). PolyP also plays a myriad of roles in fungi. For instance, in both *C. neoformans* and *Candida albicans*, polyP is involved in cation resistance and detoxification (5,32). Emerging research on cell cycle progression in *Saccharomyces cerevisiae* also revealed a new role for polyP in DNA replication and dNTP synthesis (33). In *Ustilago maydis*, another basidiomycetous fungal pathogen, polyP directly influences virulence and symptom development (26,34).

A number of key proteins are involved in polyP metabolism (5,35). In *S. cerevisiae* (and other fungi), synthesis and vacuolar translocation of polyP is mediated by a polyP synthetase known as vacuolar transporter chaperone 4 or Vtc4 (5,36). Cells deficient in polyP synthetase have no detectable polyP and, for *C. neoformans,* this results in altered proliferation in the host and an impaired ability to trigger blood coagulation (37). Additionally, the genes *XPP1* and *EPP1* in *C. neoformans* encode putative exo- and endopolyphosphatases that catalyze the hydrolysis or mobilization of inorganic polyP in response to phosphate-limiting conditions (5). Cells lacking exo- and endopolyphosphatases have significantly increased levels of polyP (5). Here we demonstrate that mutants lacking Xpp1 and Epp1 display an altered cell surface architecture, defects in mitochondrial morphology and impaired growth on medium containing inositol. These mutants are also attenuated in mice, and infected animals display an altered immune response thus supporting the conclusion that polyphosphatases are important for fungal pathogenesis.

## Results

### Mutants deficient in polyP mobilization exhibit attenuated virulence

Previously, we demonstrated that impaired synthesis of polyP does not influence virulence in a mouse inhalational model (5). However, how enhanced polyP levels impact virulence in *C. neoformans* has not been studied. Given that defects in polyphosphatases reduce virulence in bacterial and other fungal pathogens (38,39), we hypothesized that the disruption of polyP mobilization and overaccumulation of intracellular polyP would also impact the virulence of *C. neoformans*. To test this hypothesis, we assessed the ability of *xpp1*Δ, *epp1*Δ, and *xpp1*Δ*epp1*Δ mutants (two independent strains of each) defective in polyP mobilization to cause disease in a mouse model of cryptococcosis. We challenged BALB/c mice by intranasal inoculation with the wild type (WT) strain (H99) or the mutant strains (*xpp1*Δ, *epp1*Δ, and *xpp1*Δ*epp1*Δ), and monitored disease progression. In contrast to the WT strain, which caused all infected mice to succumb to the disease by day 16, each of the mutants was attenuated such that the infected mice exhibited delayed disease symptoms and mortality (Fig 1A). The difference in virulence for the double mutants was also evident when disease progression was monitored by weight loss. Mice infected with the WT strain lost weight over a period of 15 days whereas the double mutant-infected mice showed a more gradual decline in weight. (Fig 1B). An examination of fungal burden in organs harvested from infected mice revealed significantly increased colony forming units (CFUs) in the livers, kidneys, spleens and brains of mice infected with the double mutants at the experimental endpoint (Fig 1C). Surprisingly, we observed no significant differences in the lung fungal burden among the strains tested.

**Fig 1.**
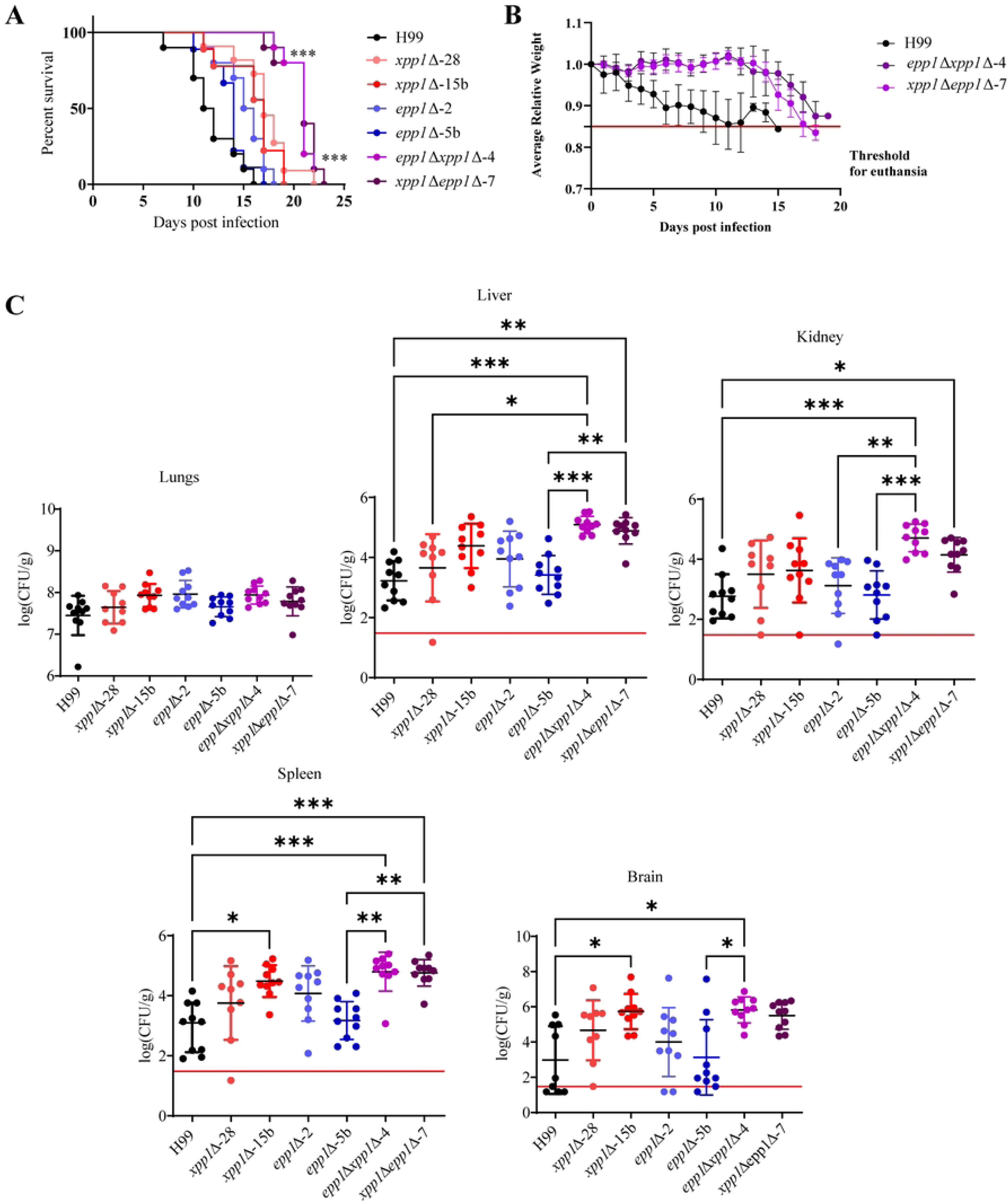
Loss of polyphosphatases influences virulence in a mouse model of cryptococcosis. **(A)** Survival of BALB/c mice intranasally inoculated with cells of WT strain (H99), two independent *xpp1*Δ mutants, two *epp1*Δ mutants, or two *xpp1*Δ*epp1*Δ mutants. **(B)** The average daily mouse weight, relative to initial weight, was used to track disease progression during the first 20 days post infection (dpi). A solid red horizontal line indicates the threshold for euthanasia at 85% of the initial weight. Error bars represent standard deviation (SD) from all surviving mice (n=10) **(C)** Colony forming units (CFUs) were measured to determine fungal burden in lungs, liver, kidney, spleen and brain for mice infected with each strain. The solid red horizontal line indicates the limit of detection for each organ. Data are presented as mean ± SD. Significance indicated as *, *P* < 0.05; **, *P* < 0.01; ***, *P* < 0.001; one-way ANOVA or log-rank test.

### Mice infected with *xpp1*Δ*epp1*Δ mutants show reduced fungal burdens and altered cytokine profiles at early infection points

We next asked whether the pattern of virulence exhibited by the *xpp1*Δ*epp1*Δ mutant at experimental endpoint was also observed at earlier times (days 7 and 14). Given the delayed disease symptoms and mortality of the *xpp1*Δ*epp1*Δ infected mice, we also wondered if mutants lacking polyP degrading enzymes were also impaired in establishing infections early in the host. To test this, we monitored disease progression and measured fungal load. The *vtc4*Δ mutant, which lacks the gene encoding polyP polymerase and therefore has no detectable polyP, was also included in our investigation to compare the influence of both low and high intracellular polyP levels on virulence. At 7 days, mice infected with the *xpp1*Δ*epp1*Δ mutant showed significantly reduced fungal burdens in the lungs compared to the *vtc4*Δ mutant and the WT (Fig 2A). Liver fungal load was also significantly reduced for the mice infected with the double mutant compared to the WT strain. While we noted a consistent decrease in spleen fungal burden and an increase in brain fungal burden among mice infected with the double mutant compared to those infected with the WT strain, these variances did not reach statistical significance. Lung cytokine profiles showed decreased production of IL-4 in mice infected with the double mutant compared to WT infected mice and elevated production of IL-6 and IFNγ for mice infected with the *vtc4*Δ mutant compared to uninfected mice (Fig 2B). The histological findings at 7 days post infection (dpi) were consistent with our fungal burden analyses (Fig 2C).

**Fig 2.**
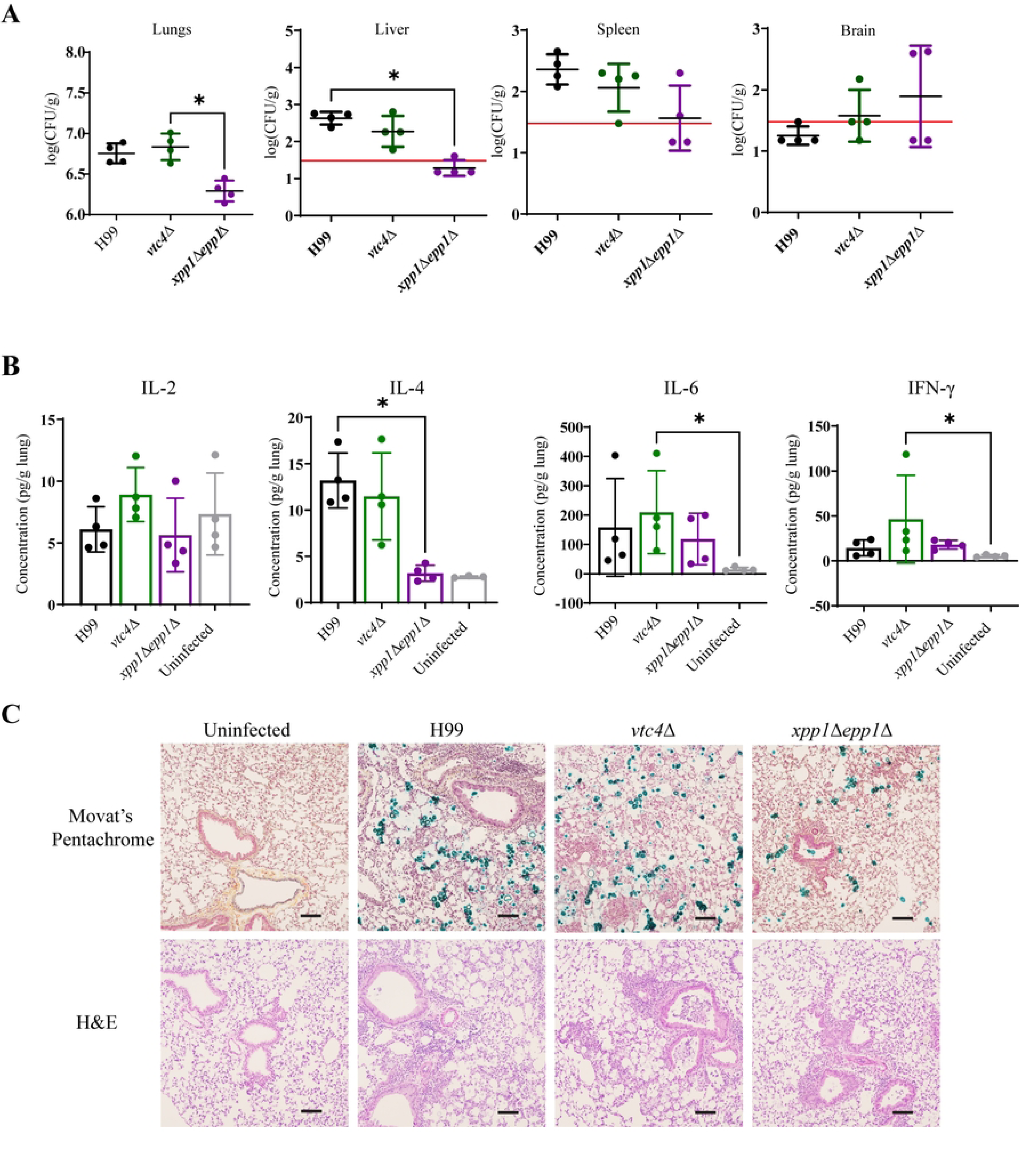
The *xpp1*Δ*epp1*Δ mutant provokes an altered immune response at day 7 and attenuated virulence in the mouse model. **(A)** Colony forming units (CFUs) were measured to determine fungal burden in lungs, liver, spleen and brain for mice infected with WT, *vtc4*Δ, or *xpp1*Δ*epp1*Δ, or treated with physiological saline (uninfected) at 7 dpi. The solid red horizontal line indicates the limit of detection for each organ. **(B)** Lung cytokine profiles of BALB/c mice infected with indicated strains or treated with physiological saline at 7 days post infection. Cytokines were extracted from lung tissue using a mixer mill and quantified using a cytometric bead array (CBA) mouse inflammation kit. **(C)** Representative histopathological micrographs of lung tissue from mice infected with WT (H99), *vtc4*Δ or *xpp1*Δ*epp1*Δ mutants, collected at 7 dpi and stained with H&E or Movat’s Pentachrome. Scale bar 100 μm. Data are presented as mean ± SD, (n = 4). Significance indicated as *, *P* < 0.05; one-way ANOVA or Mann-Whitney.

In contrast to the WT strain, mice infected with the *xpp1*Δ*epp1*Δ showed no clinical symptoms and exhibited significantly reduced fungal burdens at 14 dpi. Mice infected with the *xpp1*Δ*epp1*Δ mutant had significantly reduced fungal burden in the lung compared to all other strains tested (Fig 3A). These results are consistent with our findings at 7 dpi. We also measured cytokine production in pulmonary tissue upon infection with the strains and compared with responses uninfected. We found that mice infected with the *xpp1*Δ*epp1*Δ mutant showed significantly decreased production of IL-6 compared to WT-infected mice (Fig 3B). The histological findings at 14 dpi were consistent with our fungal burden analyses (Fig 3C). Taken together, these results suggest that a balance of polyP is required to adapt to the host environment and initiate disease. Our observations led us to hypothesize that the *xpp1*Δ*epp1*Δ mutants initially struggle to establish infection, as indicated by the low lung fungal burden observed at 7- and 14-days post-infection. This diminished fungal burden in turn stimulates weak immune responses in the BALB/c mice at earlier infection timepoints allowing the mutants to survive in the lung environment and cause disease later during infection.

**Fig 3.**
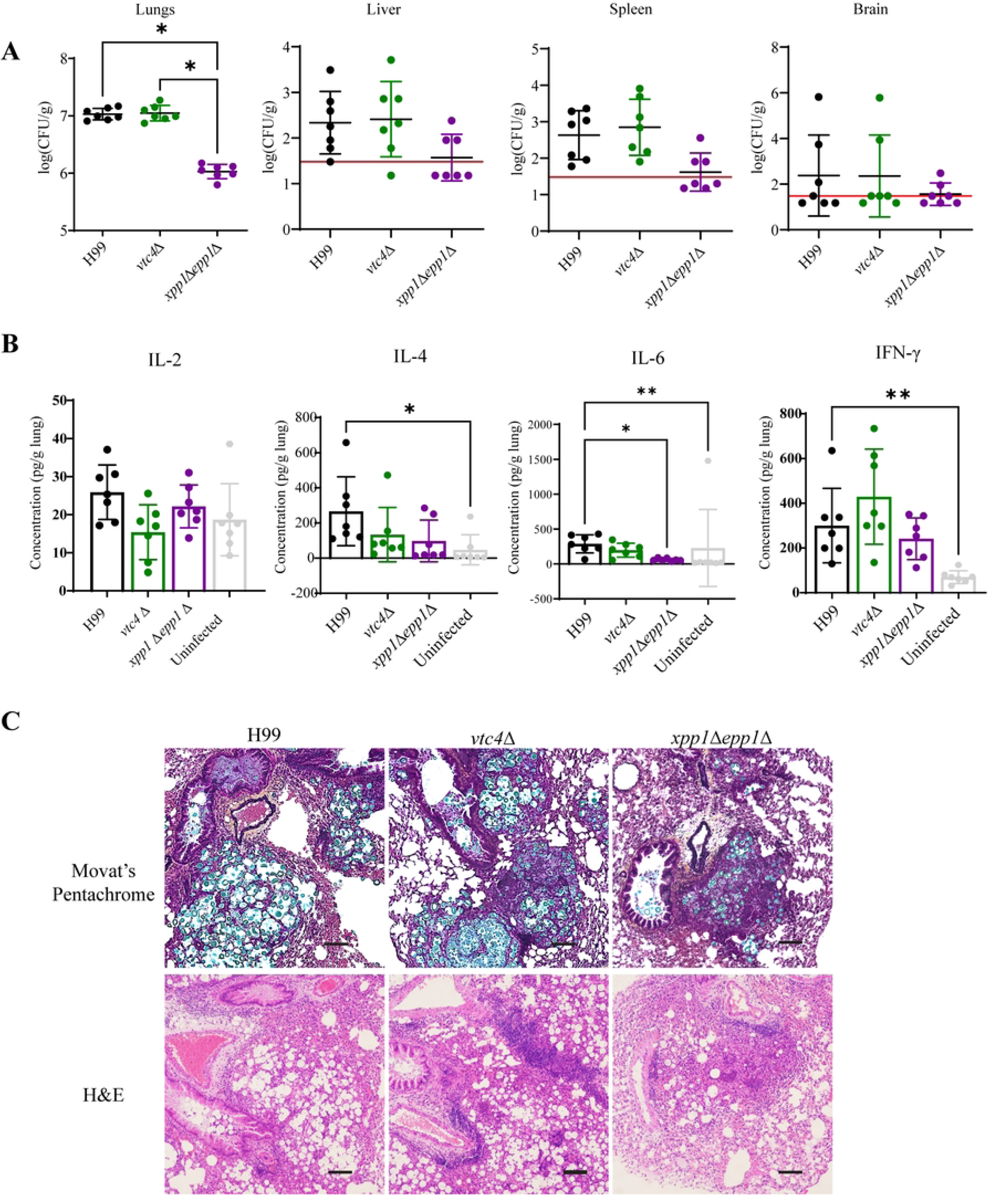
The *xpp1*Δ*epp1*Δ mutant provokes an altered immune response at day 14 and attenuated virulence in the mouse model. **(A)** Colony forming units (CFUs) were measured to determine fungal burden in lungs, liver, spleen and brain for mice infected with WT, *vtc4*Δ, or *xpp1*Δ*epp1*Δ, or treated with physiological saline at 14 dpi. The solid red horizontal line indicates the limit of detection for each organ. **(B)** Lung cytokine profiles of BALB/c mice infected with indicated strains or treated with physiological saline at 14 days post infection. Cytokines were extracted from lung tissue using a mixer mill and quantified using a cytometric bead array (CBA) mouse inflammation kit. **(C)** Representative histopathological micrographs of lung tissue from mice infected with WT, *vtc4*Δ or *xpp1*Δ*epp1*Δ mutants, collected at 14 dpi and stained with H&E or Movat’s pentachrome. Scale bar: 100 μm. Data are presented as mean ± SD. Significance indicated as *, *P* < 0.05 **, *P* < 0.01; one-way ANOVA or Mann-Whitney.

### Mice infected with the *xpp1*Δ*epp1*Δ mutant display an altered immune response

We next investigated the virulence of the *xpp1*Δ*epp1*Δ mutant by examining immune cell populations in the lungs of infected mice and mice treated with physiological saline as a control at 7 dpi by flow cytometry (Fig 4A). The lungs of the BALB/c mice infected with the double mutant had significantly reduced numbers of CD45+ cells and diminished lymphoid populations compared with those infected with the WT strain (Fig 4B). In particular, there were decreased numbers of both CD4+ and CD8+ T cells, but no differences in B cell numbers were observed (Figs 4C, S1A). There was also a significant reduction in the number of innate immune cells such as neutrophils and eosinophils (Fig 4D). The mice infected with the double mutant also exhibited decreased numbers of monocyte-derived macrophages but not monocytes (Fig 4D). Compared to the mice infected with the *vtc4*Δ mutant, infection with the *xpp1*Δ*epp1*Δ mutant resulted in reduced numbers of alveolar macrophages, NK and NKT cells in the lung (S1B and S1D Figs). These mice also had diminished type 2 conventional dendritic cells (cDC2) compared to WT infected mice in the lung (S1C Fig). We also investigated the bronchoalveolar lavage (BAL) fluid of infected mice and mice treated with physiological saline as a control at 7 dpi. Mice infected with the *vtc4*Δ mutant also showed significantly elevated levels of CD8+ T cells in the BAL fluid but we did not observe a difference in the numbers of neutrophils, eosinophils, monocytes or monocyte-derived macrophages (Fig 5A-5C). Overall, these results suggest that the mutants deficient in polyP mobilization stimulate a weaker immune response compared to infection with the WT and *vtc4*Δ strains which, in turn, might delay or prevent clearance of the double mutant from infected tissue.

**Fig 4.**
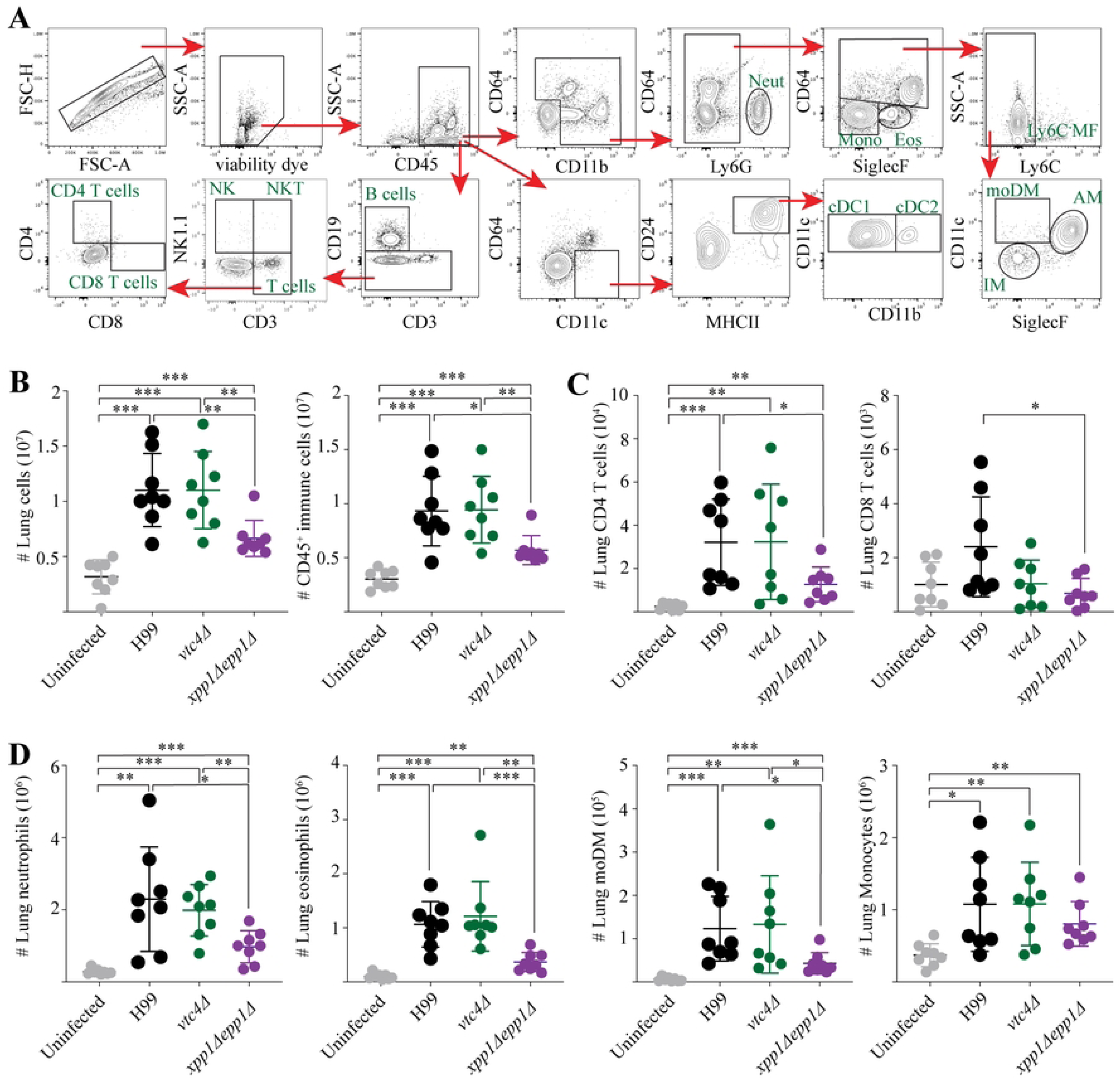
The *xpp1*Δ*epp1*Δ mutant provokes an altered immune response characterized by significantly reduced immune cell populations in lung tissue. Immune cell analysis in the lung tissue of BALB/c mice infected with WT, *vtc4*Δ, or *xpp1*Δ*epp1*Δ, or treated with physiological saline at 7 dpi. **(A)** Gating strategy, representative plots from a saline-treated lung. Doublets and debris were excluded using FSC and SSC. B cells were identified as viable CD45^+^CD19^+^CD3^-^ cells. T cells were classified as viable CD45^+^CD19^-^CD3^+^ cells and further separated as CD4+ and CD8+ T cells based on CD4^+^CD8^-^ and CD8^+^CD4^-^ respectively. Natural killer (NK) cells were gated as viable CD45^+^CD19^-^CD3^-^NK1.1^+^ cells. Natural killer T (NKT) cells were defined as viable CD45^+^CD19^-^CD3^+^NK1.1^+^ cells. Conventional dendritic cells type 1 (cDC1) and type 2 (cDC2) were identified as viable CD45^+^CD24^+^MHCII^hi^CD11c^hi^CD11b^-^ and CD11b^+^ respectively. Neutrophils (Neut) were classified as viable CD45^+^CD11b^+^CD64^-^Ly6G^+^ cells. Eosinophils (Eos) were defined as viable CD45^+^CD11b^+^CD64^-^Ly6G^-^SiglecF^+^ cells. Monocytes (Mono) were gated as viable CD45^+^Ly6G^-^CD11b^+^SiglecF^-^CD64^-^ cells. Ly6C^-^CD64^+^ macrophages (Ly6C^-^MF) were identified as viable CD45^+^Ly6G^-^CD11b^lo/+^CD64^+^Ly6C^-^ cells. Subsequently, interstitial macrophages (IM) were identified as viable CD45^+^Ly6G^-^CD11b^lo/+^CD64^+^Ly6C^-^SiglecF^-^CD11c^-^ cells. Monocyte-derived macrophages (moDM) were classified as viable CD45^+^Ly6G^-^CD11b^lo/+^CD64^+^Ly6C^-^SiglecF^-/lo^CD11c^+^ cells. Alveolar macrophages (AM) were defined as viable CD45^+^Ly6G^-^CD11b^lo/+^CD64^+^Ly6C^-^SiglecF^+^CD11c^+^ cells. **(B)** Total number of lung cells and CD45^+^ leukocytes, **(C)** adaptive CD4 and CD8 T cells, and **(D)** innate immune cells, including neutrophils, eosinophils, moDM and monocytes in the lungs. Data are presented as mean ± SD and representative of at least 2 independent experiments for each time point (n = 4 mice/time point). Significance indicated as *, *P* < 0.05; **, *P* < 0.01; ***, *P* < 0.001; unpaired Student *t* test.

**Fig 5.**
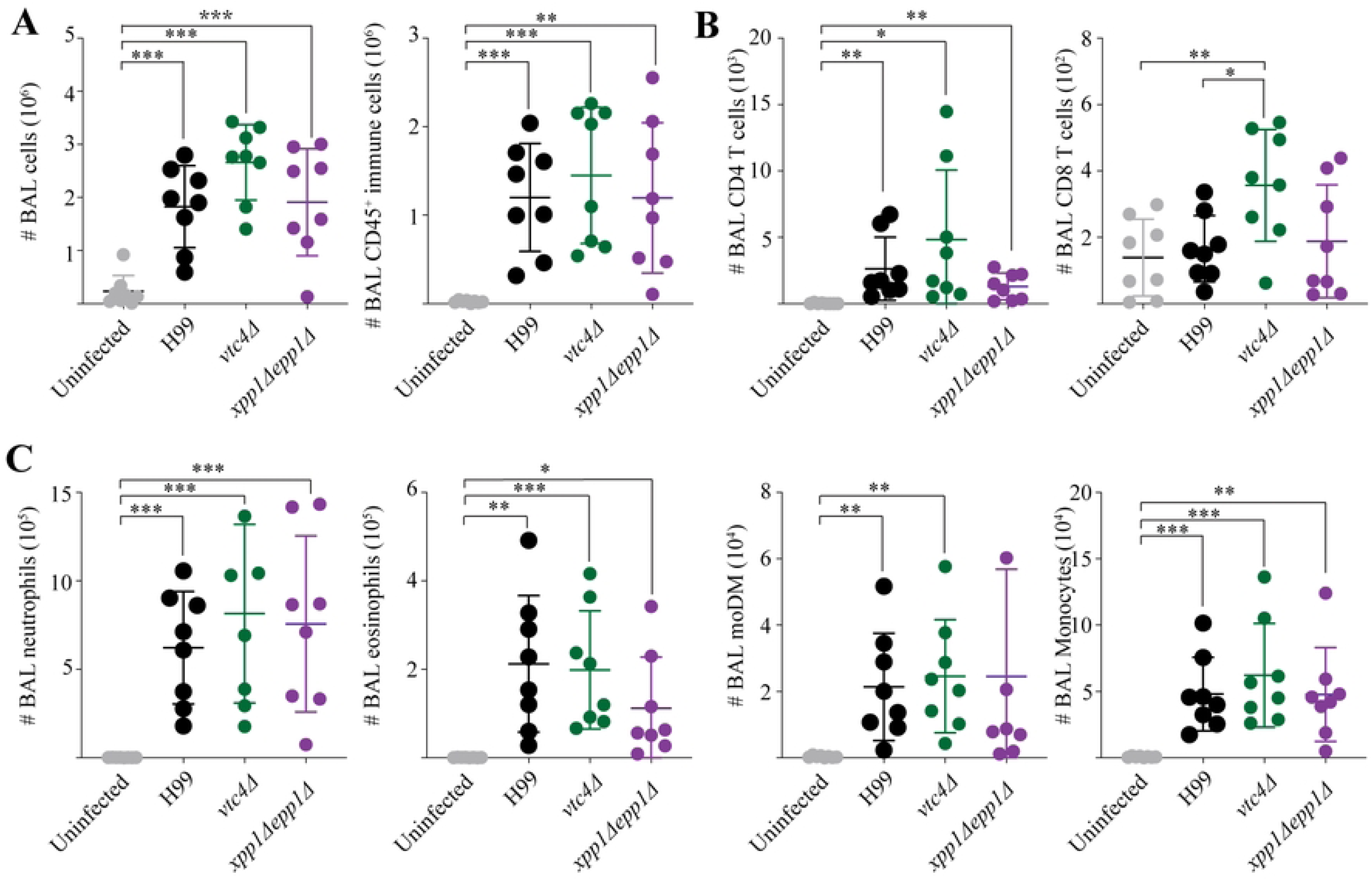
The *xpp1*Δ*epp1*Δ mutant does not cause a reduction in immune cell numbers in the BAL fluid. Immune cell analysis of bronchoalveolar lavage (BAL) fluid of BALB/c mice infected with WT, *vtc4*Δ, or *xpp1*Δ*epp1*Δ, or treated with physiological saline at 7 dpi. Gating strategy is outlined in Fig. 4A. **(A)** Total number of lung cells and CD45^+^ leukocytes, **(B)** adaptive CD4+ and CD8+ T cells, and **(C)** innate immune cells, including neutrophils, eosinophils, moDMs and monocytes in the BAL fluid. Data are presented as mean ± SD and representative of at least 2 independent experiments for each time point (n = 4 mice/time point). Significance indicated as *, *P* < 0.05; **, *P* < 0.01; ***, *P* < 0.001; unpaired Student *t* test.

### Mutants defective in polyP synthesis and mobilization are attenuated for virulence

We also constructed two independent triple deletion mutants lacking the *vtc4*, *xpp1*Δ and *epp1*Δ genes (designated *vtc4*Δ*epp1*Δ*xpp1-14 and vtc4*Δ*epp1*Δ*xpp1*-27j) to examine the impact of defective polyP synthesis and mobilization. Similar to the *vtc4*Δ mutant, the triple deletion mutants exhibit impaired polyP accumulation (characterized by the absence of detectable polyP) and enhanced susceptibility to zinc (S2 Fig). We challenged mice by intranasal inoculation with the WT strain or two independent triple deletion mutants and monitored disease progression. In contrast to the WT strain, which caused all infected mice to succumb to the disease by day 16, each of the triple mutants was attenuated in our model, and the mice exhibited delayed disease symptoms and mortality with the last infected mouse euthanized at day 26 (Fig 6A). Compared to the *epp1*Δ *xpp1*Δ double mutants, the virulence of the triple mutants was relatively more attenuated. There was a 3-day difference in the time of mortality onset between the two groups: 50% of *epp1*Δ*xpp1*Δ-infected mice succumbed to the infection by day 21, whereas 50% of mice infected with the triple mutants succumbed to the infection by day 24 (Fig 1A and 6A). Similar to mice infected with the double mutants, mice infected with the *vtc4*Δ*epp1*Δ*xpp1*Δ mutants showed a more gradual decline in weight loss (Fig 6B). Moreover, mice infected with the triple mutants exhibited significantly increased fungal burdens in the lungs, livers, kidneys, spleens, and brains at the experimental endpoint (Fig 6C). While mice infected with the WT strain consistently lost weight post infection, mice infected with both mutants defective in polyP mobilization experienced a brief recovery period and weight gain before succumbing to the infection at later timepoints (Fig 1B and 6B). Taken together, these results suggest that these mutants are slower to proliferate in mice and show greater dissemination to other organs compared to WT strain indicating contributions of the endo- and exopolyphosphatases to virulence in BALB/c mice.

**Fig 6.**
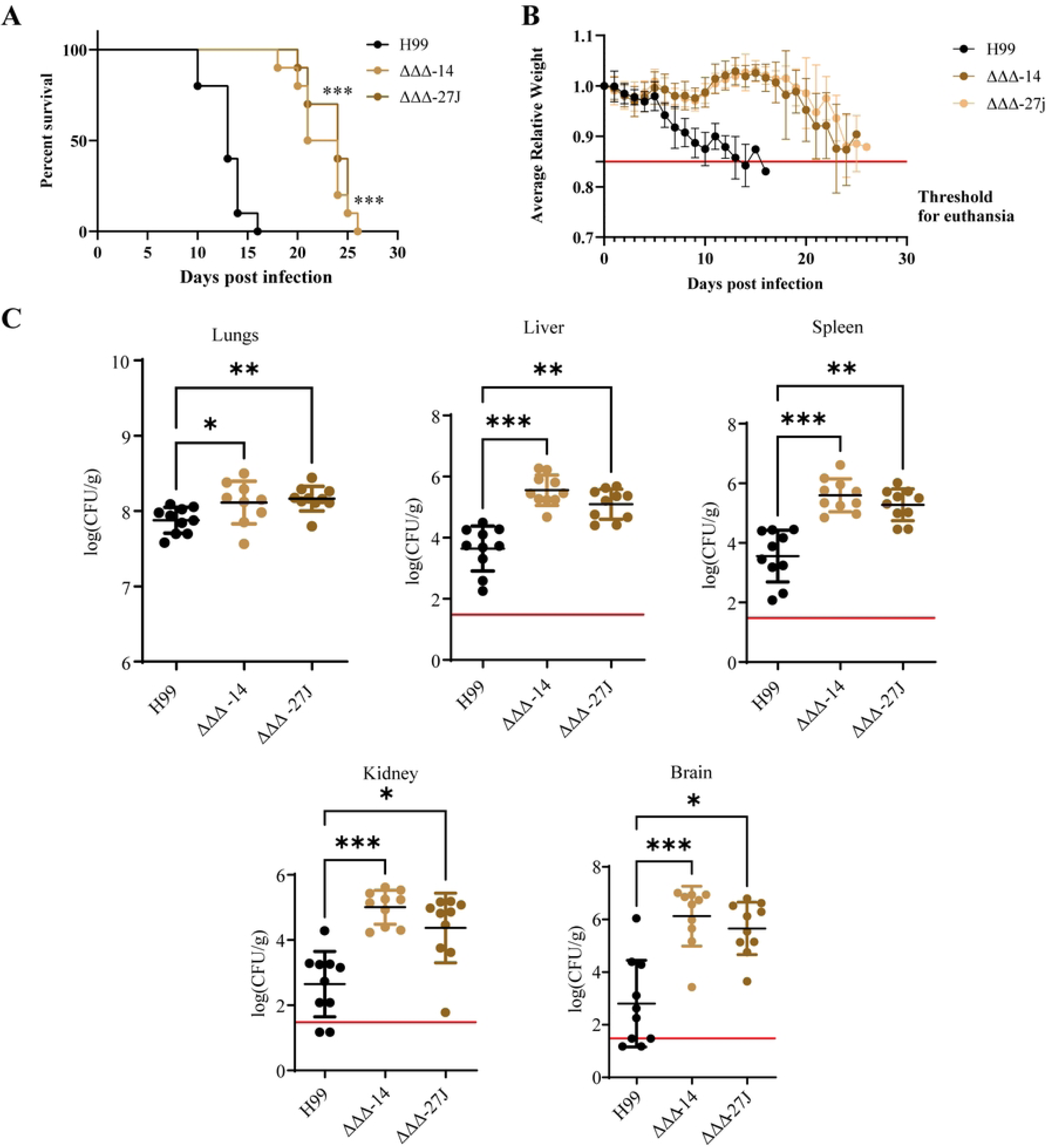
Loss of polyP synthesis and degradation enzymes impacts virulence in a mouse model of cryptococcosis. **(A)** Survival of BALB/c mice intranasally inoculated with cells of the WT strain or two independent *vtc4*Δ*epp1*Δ*xpp1*Δ (ΔΔΔ) mutants. **(B)** The average daily mouse weight, relative to initial weight, was used to track disease progression during the first 28 days post-infection. A solid red horizontal line indicates the threshold for euthanasia at 85% of the initial weight. Error bars represent standard error of means (SEM) from all surviving mice (n = 10). **(C)** Colony forming units (CFUs) were measured to determine fungal burden in lungs, liver, kidney, spleen and brain for mice infected with each strain. The solid red horizontal line indicates the limit of detection for each organ. Data are presented as mean ± SD. Significance indicated as *, *P* < 0.05; **, *P* < 0.01; ***, *P* < 0.001; one-way ANOVA or log-rank test.

### Impaired polyP mobilization is linked to cell size *in vivo*

Given that phosphate acts as a signal for small cell induction and that decreased cell size is conducive for extrapulmonary dissemination, we hypothesized that the elevated dissemination phenotype observed in mice infected with the double and triple deletion mutants is due to smaller yeast cell size *in vivo* (40). Indeed, our investigation of *C. neoformans* cells isolated from lung homogenates revealed a connection between impaired polyP breakdown and cell size. We discovered that cells defective in polyP mobilization were significantly smaller in diameter compared to WT cells *in vivo* (Fig 7A-7C). Mice infected with the *xpp1*Δ*epp1*Δ mutants also exhibited disrupted lung architecture at endpoint (Fig 7D). Movat’s Pentachrome staining also revealed an enhanced accumulation of mucin in the lung airways of mice infected with the *xpp1*Δ*epp1*Δ mutants (Fig 7D). Similar to mice infected with the *xpp1Δepp1Δ* mutants, mice infected with the triple mutants also showed disrupted lung architecture, and we noted that the lungs from mice infected with mutants impaired in polyP mobilization were enlarged (Fig 7C) and difficult to homogenize.

**Fig 7.**
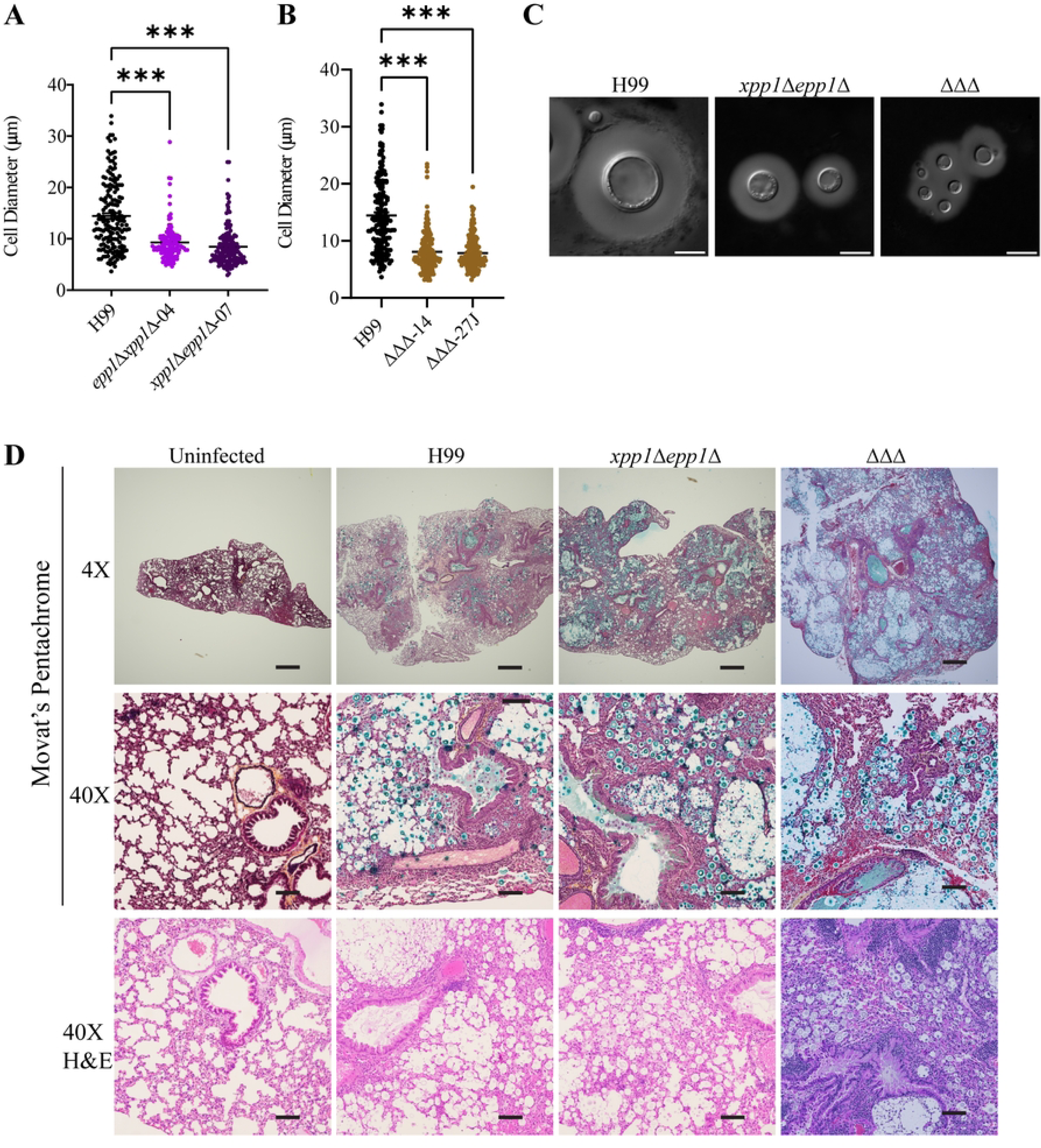
Defects in polyP mobilization results in smaller cell diameters and enhanced disruption of murine lung architecture. Yeast cells of the WT strain, two independent *xpp1*Δ*epp1*Δ mutants or two *vtc4*Δ*epp1*Δ*xpp1*Δ (ΔΔΔ) mutants were isolated from the lungs of infected mice (n = 10) and **(C)** stained with India ink to visualize polysaccharide capsules. (**A, B**) Cell size was measured manually and at least 100 cells were analyzed per strain. Representative differential interference contrast (DIC) micrographs are shown. Scale bar 10 μm. **(D)** Representative histopathological micrographs of lung tissue from mice infected with WT, *xpp1*Δ*epp1*Δ or ΔΔΔ mutants, collected at experimental endpoint and stained with H&E or Movat’s Pentachrome. Scale bar: 4X = 500 μm; 40X = 100 μm. Data are presented as mean ± SEM. Significance indicated as ***, *P* < 0.001; Mann-Whitney.

### Mutants defective in polyP synthesis and mobilization are impaired for uptake and survival in macrophages

Given the protective roles of polyP against the harsh environment of the host phagolysosome, we next examined the interactions of the mutants with murine macrophages by assessing uptake and survival of opsonized yeast cells. We found that the mutants defective in synthesis and mobilization of polyP exhibited significantly reduced uptake by bone-marrow derived primary macrophages (BMDMs) (Fig 8A and 8B). That is, the number of internalized yeast cells 2 hours post infection was significantly reduced for the *vtc4*Δ, *xpp1*Δ*epp1*Δ, and *vtc4*Δ*epp1*Δ*xpp1*Δ mutant strains compared to the WT strain. However, only mutants defective in polyP mobilization, i.e., *xpp1*Δ*epp1*Δ and *vtc4*Δ*epp1*Δ*xpp1*Δ, showed reduced survival in the primary macrophages (Fig 8B and 8C). In other words, the number of internalized yeast cells 24 hours post infection was significantly reduced for mutants lacking polyphosphatases but not for the *vtc4*Δ mutant. We confirmed these findings independently by counting colony forming units (CFUs) after macrophage lysis (Fig 8D). We also assessed uptake of the mutants by J774A.1 murine macrophage-like cell line. Consistent with our findings with BMDMs, mutants deficient in synthesis and mobilization of polyP exhibited reduced uptake by J774A.1 macrophages (Fig 8E and 8F). Overall, these results suggest that a balance in polyP levels is crucial for uptake by and survival inside macrophages.

**Fig 8.**
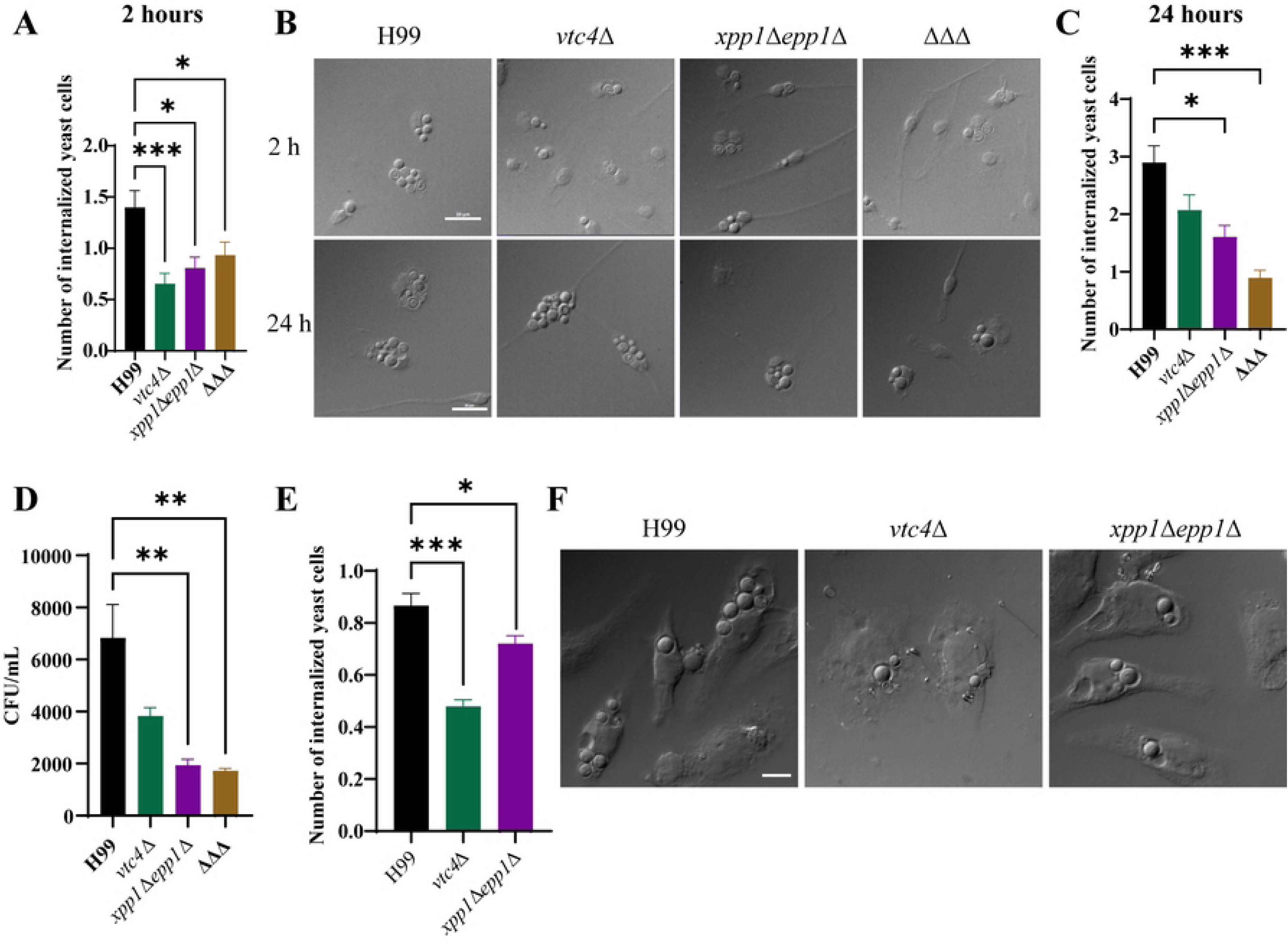
Defects in polyP metabolism reduce the uptake and survival of *C. neoformans* in murine macrophages. **(A)** Uptake of opsonized WT, *vtc4*Δ, *xpp1*Δ*epp1*Δ, or *vtc4*Δ*epp1*Δ*xpp1*Δ (ΔΔΔ) strain in bone marrow derived macrophages is shown after 2 hours of interaction. **(B)** Representative DIC micrographs of bone-marrow derived primary macrophages (BMDMs) with internalized yeast cells at 2 hours and 24 hours of interaction are shown. Scale bar 30 μm. **(C)** Survival of opsonized WT, *vtc4*Δ, *xpp1*Δ*epp1*Δ, or *vtc4*Δ*epp1*Δ*xpp1*Δ (ΔΔΔ) strain in bone marrow derived macrophages is shown after 24 hours of interaction. **(D)** Survival of opsonized WT, *vtc4*Δ, *xpp1*Δ*epp1*Δ, or *vtc4*Δ*epp1*Δ*xpp1*Δ in bone marrow derived macrophages after 24 hours of interaction measured by counting colony forming units. **(E, F)** Uptake of opsonized WT, *vtc4*Δ, or *xpp1*Δ*epp1*Δ strain in J774A.1 macrophage-like cell line is shown after 2 hours of interaction. Scale bar 10 μm. Number of internalized *C. neoformans* cells per macrophage were counted by analyzing 500 macrophages per strain for each experiment. Data are presented as mean ± SEM and representative of at least 3 independent experiments. Significance indicated as *, *P* < 0.05; **, *P* < 0.01; ***, *P* < 0.001; Kruskal-Wallis nonparametric test.

### Loss of Xpp1 and Epp1 alters cell surface architecture and influences the elaboration of virulence factors

Previous studies revealed the presence of polyP in cell wall and capsular compartments for *C. neoformans* (41). That is, polyP is required for correct capsular assembly and cells defective in polyP synthesis show an altered capsular architecture characterized by thicker polysaccharide fibers (41). Given the importance of phosphate and polyP in polysaccharide capsule assembly and elaboration, we hypothesized that mutants accumulating elevated polyP due to defects in the polyP degrading enzymes (Epp1 and Xpp1) would exhibit differences in cell wall and capsular architecture. To test this hypothesis, we investigated the cell surface by India ink staining and scanning electron microscopy (SEM) and found cell surface architectural changes for mutants defective in polyP synthesis and mobilization. In particular, the *xpp1*Δ*epp1*Δ mutant was defective in elaboration of the polysaccharide capsule and capsules produced by this mutant were smaller compared to the WT and *vtc4*Δ strains (Fig 9A and 9B). It was interesting that despite a reduction in the size of the polysaccharide capsule, the double mutant showed reduced uptake in the macrophages. SEM analysis of the capsular ultrastructure revealed that the double mutant displayed capsular fibers that were more spread apart and scattered compared to WT and *vtc4*Δ mutant cells (Fig 9A). The *vtc4*Δ*epp1*Δ*xpp1*Δ mutants also elaborated smaller capsules compared to the WT and the *vtc4*Δ mutant (Fig 9B). These mutants also showed perturbed capsular ultrastructure (Fig 9A).

**Fig 9.**
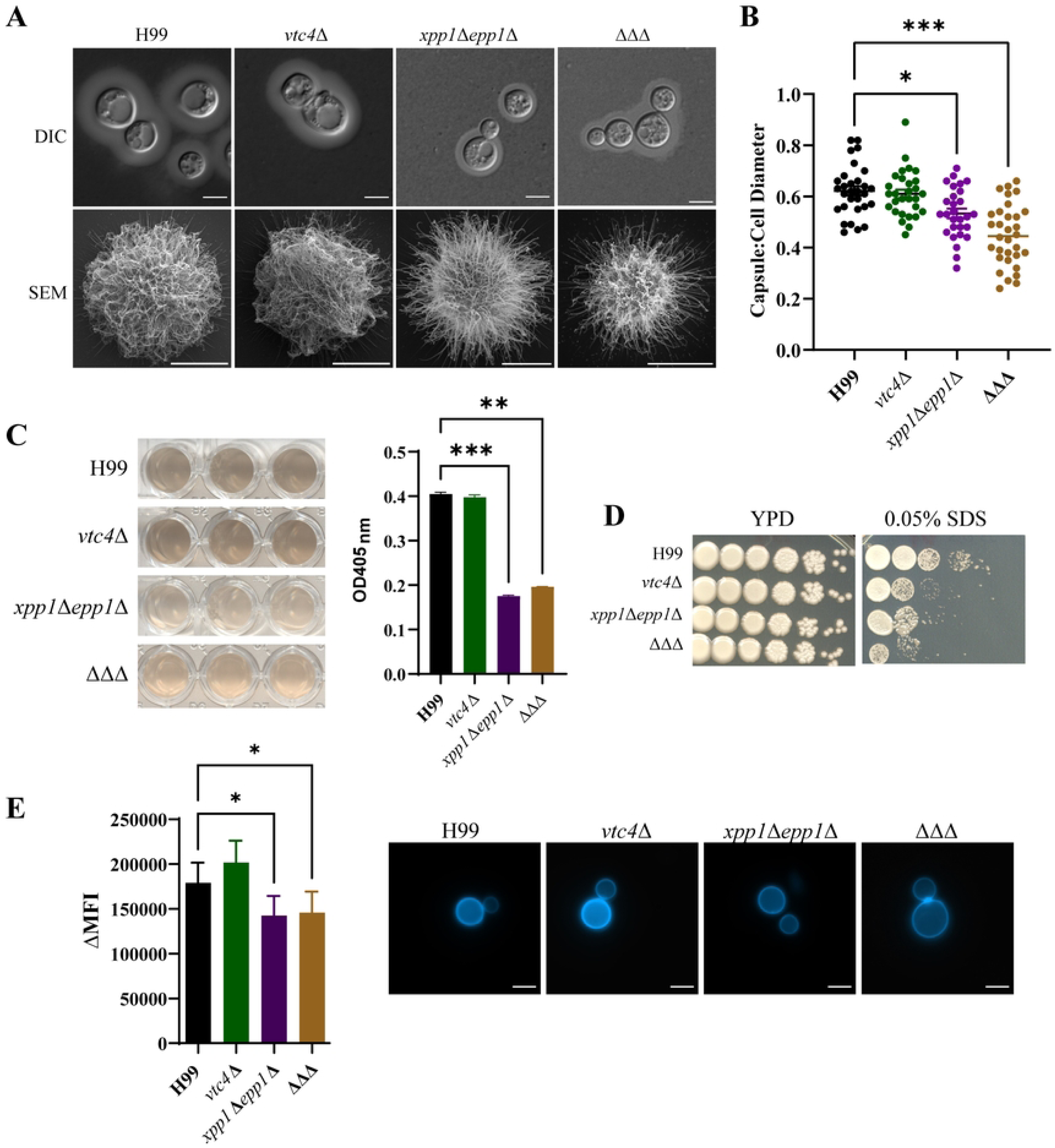
Mutants deficient in polyP mobilization have altered cell surface architecture. **(A)** WT, *vtc4*Δ, *xpp1*Δ*epp1*Δ or *vtc4*Δ*epp1*Δ*xpp1*Δ strains were inoculated into capsule inducing medium (CIM) and incubated at 30°C for 48 hours. Capsule formation and structure were assessed by India ink staining or scanning electron microscopy (SEM) for the indicated strains. **(B)** Ratio of capsule thickness to cell diameter was quantified for at least 50 cells per strain. **(C)** WT, *vtc4*Δ, *xpp1*Δ*epp1*Δ or *vtc4*Δ*epp1*Δ*xpp1*Δ strains were inoculated into L-DOPA liquid medium and incubated at 30°C for 48 hours. Melanin production was quantified using OD405 values. **(D)** Indicated strains were serially diluted and spotted onto solid YPD agar with or without 0.05% SDS. The plates were then incubated at 30 °C for 5 days before being photographed. **(E)** WT, *vtc4*Δ, *xpp1*Δ*epp1*Δ or *vtc4*Δ*epp1*Δ*xpp1*Δ strains were stained with calcofluor white (CFW, 100 μg/mL) to visualize chitin. Chitin content was quantified using flow cytometry and analyzed by FlowJo. ΔMFI = change in mean fluorescence intensity. Scale bar 5 μm. Data are presented as mean ± SEM and representative of at least 3 independent experiments. Significance indicated as *, *P* < 0.05; **, *P* < 0.01; ***, *P* < 0.001; one-way ANOVA.

The attenuated virulence of the *xpp1*Δ*epp1*Δ and *vtc4*Δ*epp1*Δ*xpp1*Δ mutants in the murine inhalational model and reduced survival in murine macrophages also prompted us to hypothesize that the polyphosphatases, Xpp1 and Epp1 may also influence other major virulence factors. We therefore assessed the *xpp1*Δ*epp1*Δ mutant for growth at elevated temperatures, melanin production, and elaboration of the polysaccharide capsule. We discovered the double mutants displayed normal growth at 39°C and that loss of both Xpp1 and Epp1 resulted in reduced production of melanin on medium containing L-DOPA as a substrate (Fig 9C). The altered cell surface also prompted us to test the ability of these mutants to respond to membrane and cell wall stress agents such as SDS, Congo Red, CFW and caffeine. We discovered that mutants defective in both polyP synthesis and mobilization are sensitive to SDS indicating a role of polyP in membrane integrity (Fig 9D). Surprisingly, all of the mutants grew normally on YPD supplemented with Congo Red, CFW and caffeine (S3 Fig). However, staining with calcofluor white (CFW) revealed reduced chitin content for both the *xpp1*Δ*epp1*Δ and *vtc4*Δ*epp1*Δ*xpp1*Δ mutants compared to WT and the *vtc4*Δ mutant (Fig 9E).

### Xpp1 and Epp1 influence mitochondrial morphology and stress resistance

A recent study linking mitochondrial inorganic polyP to the generation of reactive oxygen species (ROS) in mammalian cells prompted us to analyze ROS and mitochondria-related phenotypes for our mutants (42). We first assessed accumulation of intracellular ROS using 2ʹ,7ʹ-Dichlorofluorescein Diacetate (DCFDA). Both *xpp1*Δ*epp1*Δ and *vtc4*Δ*epp1*Δ*xpp1*Δ mutants showed elevated levels of intracellular ROS compared to WT (Fig 10A and 10B). The *vtc4*Δ mutant showed a reduction of intracellular ROS levels. We next assessed the ability of the mutants to respond to ROS stress agents such as hydrogen peroxide (H_2_O_2_), *tert*-Butyl hydroperoxide (tBuOOH), paraquat and plumbagin. We discovered that mutants defective in polyP mobilization were sensitive to H_2_O_2_ at elevated temperatures (Fig 10C). The *vtc4*Δ mutant, on the other hand, displayed slight resistance to H_2_O_2_ at elevated temperatures (Fig 10C). All of the mutants grew normally on media supplemented with other ROS stress agents (S4A Fig).

**Fig 10.**
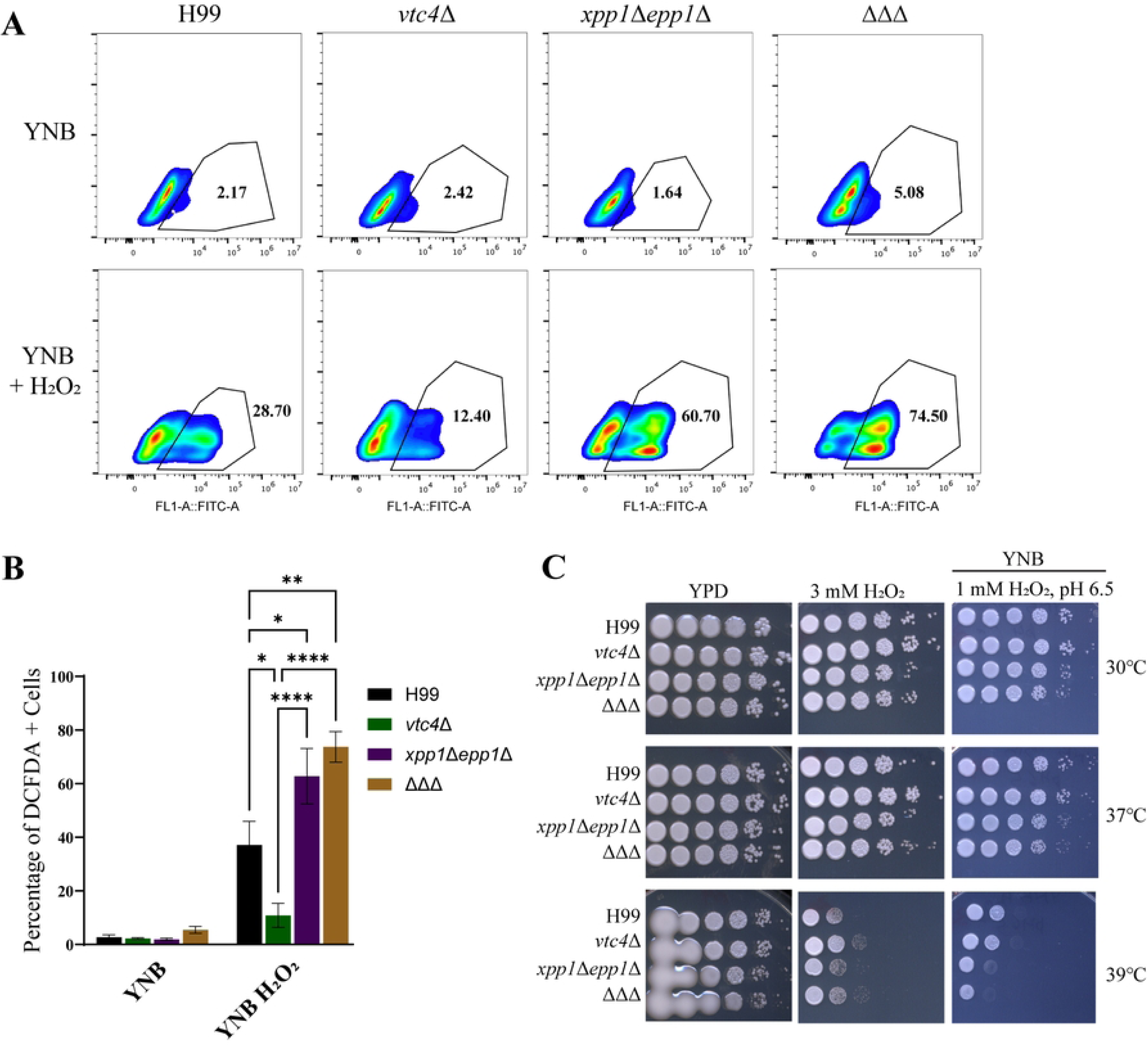
Defects in polyP mobilization confer sensitivity to reactive oxygen species stress. (A-B) Flow cytometry analysis of intracellular ROS levels for WT, *vtc4*Δ, *xpp1*Δ*epp1*Δ, or *vtc4*Δ*epp1*Δ*xpp1*Δ (ΔΔΔ) strains. Indicated strains were grown in YPD overnight at 30°C, transferred to yeast nitrogen base (YNB) liquid medium for 3-4 hours and log cells were then treated with or without H_2_O_2_ (1mM) for 1 hour at 30°C. ROS levels were measured by staining the cells with ROS indicator dye 2ʹ,7ʹ-Dichlorofluorescein Diacetate (DCFDA, 16 μM) for 1 hour. **(C)** Indicated strains were serially diluted and spotted onto solid YPD or YNB agar with or without H_2_O_2_. The plates were then incubated at 30°C, 37°C or 39°C for 2-4 days before being photographed. Data are presented as mean ± SEM and representative of at least 4 independent experiments. Significance indicated as *, *P* < 0.05; **, *P* < 0.01; ***, *P* < 0.001; ****, *P* < 0.0001; one-way ANOVA.

Previous studies have highlighted the significance of mitochondria in fungal pathogens’ adaptation to diverse stress conditions (43). Mitochondrial morphology, for example, has been linked to the response to host conditions during *C. neoformans* infections (44,45). In light of the observed sensitivities of the polyP mobilization defective mutants under oxidative stress condition (Fig 10), we focused on examining potential changes in mitochondrial morphology in the deletion mutants following exposure to oxidative stress. To investigate how oxidative stress impacts mitochondrial morphology in polyP mobilization defective mutants, we employed both MitoTracker Orange and nonyl acridine orange (NAO). These dyes allowed us to visualize both active mitochondria, based on membrane potential, and mitochondria independent of their membrane potential. We found that loss of both Xpp1 and Epp1 caused altered morphology compared to the WT strain. More specifically, the *xpp1*Δ*epp1*Δ and *vtc4*Δ*epp1*Δ*xpp1*Δ mutants exhibited reduced number of active mitochondria compared WT or *vtc4*Δ mutant cells regardless of the ROS stress conditions. (Fig 11A). This result prompted an assessment of the ability of the mutants to respond to mitochondrial stress provoked by growth on media supplemented with the electron transport chain inhibitors rotenone (complex I), antimycin A, myxothiazol (complex III), SHAM (alternative cytochrome oxidative pathway), or KCN (complex IV). Notably, mutants defective in polyP mobilization were sensitive to SHAM and KCN but grew normally on media supplemented with antimycin A, myxothiazol and rotenone (Figs 11B and S4B). To further determine whether the defects in mitochondrial morphology were linked to altered function, we evaluated alternative carbon source utilization by WT and *xpp1*Δ*epp1Δ* mutants. While the *xpp1*Δ*epp1*Δ mutants showed no difference in growth on YNB liquid media supplemented with glucose or acetic acid, there was a significant growth defect on YNB liquid media supplemented with inositol (Figs 11C and S4C).

**Fig 11.**
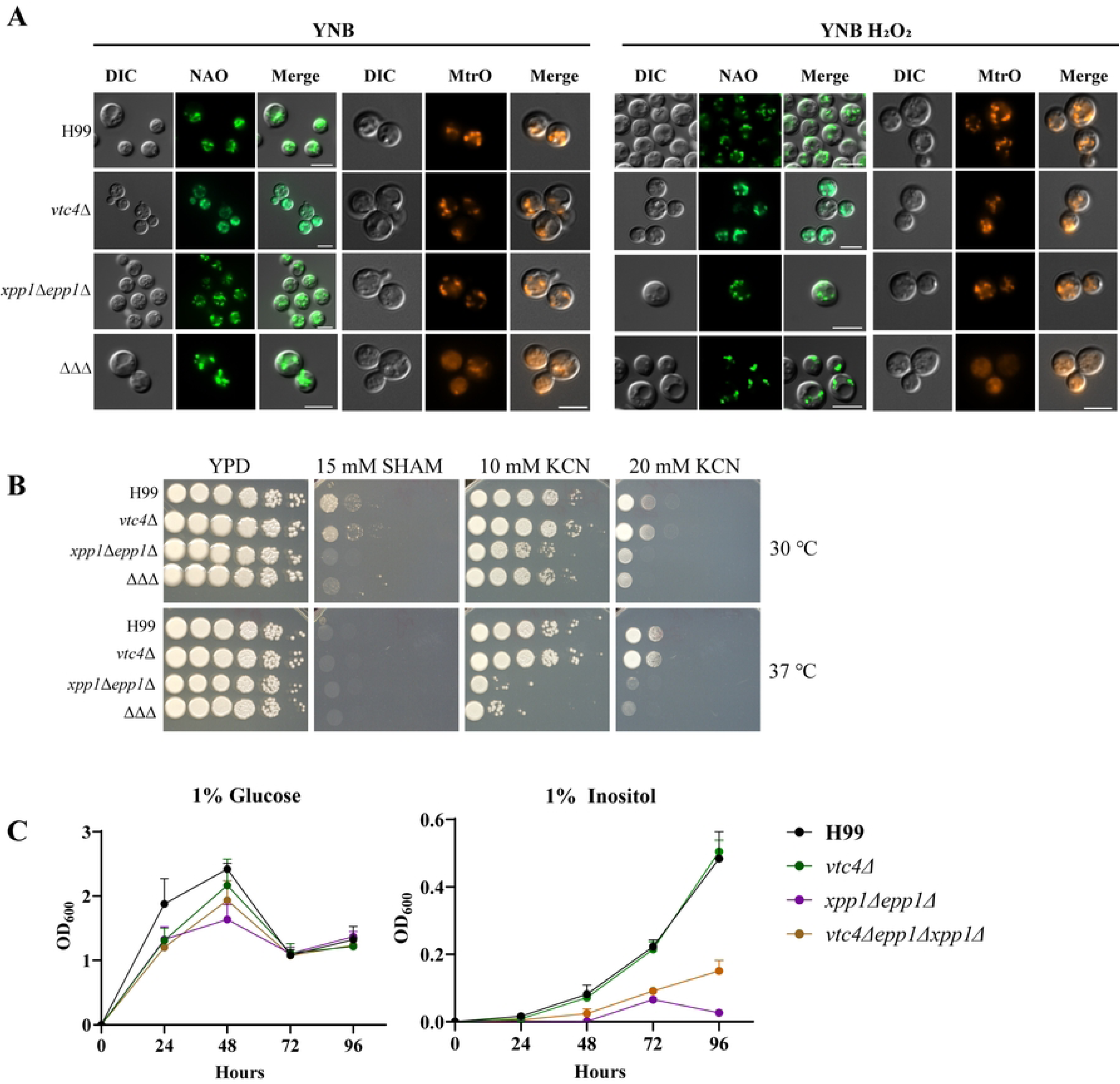
Defects in polyP mobilization impact mitochondrial morphology and impact growth on media supplemented with 2% inositol. **(A)** Representative widefield fluorescence micrographs showing mitochondrial morphologies in WT, *vtc4*Δ, *xpp1*Δ*epp1*Δ, or *vtc4*Δ*epp1*Δ*xpp1*Δ strains. Indicated strains were stained with the mitochondria specific dyes, MitoTracker Orange CMTMRos (200 nM) or nonyl acridine orange (NAO, 200 ng/μL) for 30 minutes. Scale bar 5 μm. Data representative of three independent experiments representing at least 100 cells per treatment. **(B)** Indicated strains were serially diluted and spotted onto solid YPD agar with or without mitochondrial stress agents, SHAM and KCN. The plates were then incubated at 30°C for 2-4 days before being photographed. **(C)** Liquid growth assays on glucose and inositol for the indicated strains. *C. neoformans* yeast cells were grown in YNB without amino acids with 1% glucose or 1% inositol in test tubes containing 5 mL of the indicated growth medium. The tubes were incubated at 30°C, 200 rpm for 120 hours and optical densities (OD600) were measured every 24 hours. Data are presented as mean ± SD and representative of at least 3 independent experiments.

## Discussion

In this study, we have improved our understanding of the contribution of polyP homeostasis in cryptococcal disease and highlighted new roles for polyphosphatases in cell surface architecture and mitochondrial functions. Previous studies have revealed that loss of either Xpp1 or Epp1 results in polyP accumulation along with an acquired resistance to zinc (5). In the present study, we have shown that both Xpp1 and Epp1 are required for virulence in the mouse inhalation model. We also discovered that mutants defective in polyP mobilization display sensitivity to cell membrane stress, reactive oxygen species, and ETC inhibitors. They also have decreased chitin content, reduced formation of capsule and melanin, and an increased proportion of smaller cells *in vivo*. In addition, these mutants display a defect in uptake by murine macrophages. Mutants lacking Xpp1 and Epp1 also show reduced survival in murine macrophages suggesting a role for these enzymes in survival in the macrophage phagolysosome. We also uncovered contributions of polyP mobilization to cell surface architecture. Previous studies have implicated polyP at the cell surface and shown that polyP is required for correct capsular assembly in *C. neoformans* (41). Expanding upon this finding, we found that mutants lacking Xpp1 and Epp1 exhibit abnormal polysaccharide capsular fibers, which are characterized by greater dispersion and scattered distribution compared to polysaccharide fibers elaborated by the WT or *vtc4*Δ mutant strain. This discovery implies a novel function for polyphosphatases in elaboration of the cryptococcal polysaccharide capsule.

Considerable evidence suggests that polyP interacts with mitochondria and impacts mitochondrial and other cellular functions in a variety of ways. In mammalian cells, polyP co-localizes to mitochondria and plays a crucial role in maintaining cellular bioenergetics as well as regulating oxidative phosphorylation and glycolysis (42,46,47). PolyP is also known to regulate the calcium concentration in mitochondria which in turn is needed for ATP production (47). Interestingly, we discovered that mutants with impaired polyP mobilization were only sensitive to ETC stress agents, KCN and SHAM, inhibitors of cytochrome oxidase and alternative oxidase, respectively. This finding suggests that enzymes involved in polyP breakdown have alternative functions in the later part of the electron transport chain and may be more directly involved in ATP synthesis. A recent study investigating mitochondrial bioenergetics revealed that depletion of mitochondrial polyP results in increased oxidative stress conditions due to a dysregulation of the oxidative phosphorylation and these conditions are characterized by increased intracellular H_2_O_2_ levels (42). We suspect that loss of Xpp1 and Epp1 in *C. neoformans* cells leads to a similar outcome as mitochondrial polyP is locked away and is not available for regulation of redox homeostasis. This difference, coupled with perturbed mitochondrial function and morphology, might explain elevated ROS levels in *C. neoformans* mutants deficient in polyphosphatases.

With key structural and signalling roles, inositol is an important nutrient for fungal pathogens and *C. neoformans* can use inositol as a sole carbon source (48). Inositol polyphosphates (IPs) and inositol pyrophosphates (PP-IPs) are involved in regulation of diverse cellular processes such as chromatin remodeling, calcium and phosphate homeostasis, and cell cycle (37). In *Schizosaccharomyces pombe*, inositol pyrophosphate is involved in phosphate homeostasis (49). SPX domains, present on a number of Pi-associated proteins including VTC complex proteins, bind to inositol pyrophosphate and directly influence phosphate homeostasis including polyP synthesis (50). A dysregulation of IP biosynthesis in pathogenic fungi has been shown to cause cellular defects (37). Considering the significance of inositol metabolism in the cell cycle and capsule elaboration, our discovery that mutants with impaired polyP mobilization exhibit reduced capsule and decreased cell diameters *in vivo* suggests a potential inositol-mediated involvement of these enzymes in morphogenesis and virulence. Our current investigation also revealed that both *xpp1*Δ*epp1*Δ and *vtc4*Δ*epp1*Δ*xpp1*Δ mutants were incapable of utilizing inositol as an alternative carbon source. Therefore, we propose that Xpp1 and Epp1 contribute to mechanisms of regulation of inositol biosynthesis and signalling.

Our understanding of how polyP synthesis, mobilization, and virulence intersect in fungal pathogens is limited. To our knowledge, immune responses to fungal polyP have not been studied in detail. Interestingly, the influence of polyP on the immune system is quite complex and not fully understood. It is believed that polyP polymers of different chain lengths activate distinct signaling pathways and have diverse mechanisms of action (51). A recent study investigating the role of polyphosphatases in *C. albicans* revealed that polyP mobilization is required for DNA replication, and blocking polyP mobilization results in impaired cell division and formation of large pseudo hypha-like cells (38). In addition, polyP mobilization is required for the virulence of *C. albicans* (38). Emerging studies in bacterial pathogenesis have shed light on how polyP not only influences cellular functions but also modulates the host immune response. In *Mycobacterium tuberculosis,* a mutant strain lacking exopolyphosphatases is significantly reduced in the expression of virulence genes (52). Accumulation of polyP has also been shown to prevent establishment of infection in guinea pigs by *M. tuberculosis* (52). *M*. *tuberculosis* mutants defective in polyP mobilization show significantly reduced growth and survival in macrophages (52,53). High levels of bacterial polyP also serve to deter phagocytosis by macrophages and neutrophils and contribute to inhibition of phagosome acidification in phagocytes (54). PolyP can also inhibit recruitment of macrophages into tissue (55). In mammalian cells, it is hypothesized that polyP acts as signal that impacts leukocyte recruitment, function and proliferation (53,56). PolyP is also known to contribute to proinflammatory effect of activated platelets and has roles in blood coagulation (57). A recent study reported that short-chain polyP potentiates the activity of neutrophils and stimulates the release of neutrophil extracellular traps (NETs) (58). In contrast, our analysis indicates that mice infected with the *xpp1*Δ*epp1*Δ double mutant have significantly decreased numbers of CD45+ cells, neutrophils, and eosinophils in lung tissue at 7 dpi. The diminished presence of CD45+ cells in the lung may stem from an initial lower fungal burden within lung tissue, as fewer yeast cells might elicit weaker immune responses. Additionally, mice infected with the *xpp1*Δ*epp1*Δ mutant exhibit reduced IL-4 levels compared to those infected with the WT strain. IL-4 plays detrimental roles in fungal immunity by polarizing macrophages to an alternatively activated phenotype, promoting the proliferation of intracellular pathogens (59). This suggests that infection with the double mutant promotes a protective Th1-type response, potentially facilitating more effective pathogen clearance, especially at earlier time points. Overall, the *xpp1*Δ*epp1*Δ mutant stimulated a weaker immune response compared to wild type.

It is currently unclear why the mice infected with *xpp1*Δ*epp1*Δ and *vtc4*Δ*epp1*Δ*xpp1*Δ mutants ultimately succumb to the infection at later time points. It is possible that mutants with defects in polyP mobilization may struggle to establish a robust infection during earlier timepoints and may need time to adapt to the host environment. This is evidenced by mice infected with the *xpp1*Δ*epp1*Δ mutant showing notably reduced fungal burdens at day 7 and 14. This could also account for the gradual decline in murine weight and the slow but steady progression of disease in these infected mice. We speculate that the *xpp1*Δ*epp1*Δ mutant stimulates a weaker immune response characterized by reduced numbers of CD45+ cells, neutrophils, eosinophils, CD4+ T cells, CD8+ T cells in the BALB/c mice at earlier infection timepoints which allows these mutants to survive in the lung environment and cause enhanced disease later in the infection period. It is also likely that a defect in uptake by macrophages combined with a diminished presence of eosinophils permits the *xpp1*Δ*epp1*Δ to survive in the extracellular space and proliferate in the lung tissue at later infection timepoints. It is interesting that despite no differences in lung monocyte numbers between WT and double mutant strains, mice infected with the *xpp1*Δ*epp1*Δ mutant have significantly reduced monocyte-derived macrophages. This observation hints at a possible block in the differentiation of monocytes into monocyte-derived macrophages, warranting further investigation.

Here, we also demonstrated that mutants with defective mobilization of polyP are smaller in cell size *in vivo* and show enhanced extrapulmonary dissemination over time. The presence of small or ‘micro’ cells is also associated with decreased CD4 T cell counts (60) which is consistent with our results. Large cells are known to be resistant to oxidative and nitrosative stress and show reduced extrapulmonary dissemination (61). In this present study, small cell formation *in vivo* may indicate a problem with cell cycle regulation and cell division as phosphate regulation is directly linked with cell cycle in fungi. Our analyses also reveal a connection between small cell size and extrapulmonary dissemination in *C. neoformans* consistent with the findings of Denham et al. (40). A double deletion of Xpp1 and Epp1 led to increased fungal burden in liver, kidney, spleen and brain over time in infected mice.

In summary, we find that polyphosphatases, responsible for polyP mobilization, play a multifaceted role in pathogenesis with an influence on survival in macrophages and virulence in the murine model of cryptococcosis. In addition to reduced survival in macrophages, attenuated virulence may result from susceptibility to reactive oxygen species perhaps resulting from impaired mitochondrial function, and reduced formation of critical virulence factors such as capsule and melanin. These findings focus attention on polyP as an underappreciated contributor to fungal pathogenesis and modulator of the host immune response, highlighting the potential therapeutic value of polyP.

## Materials and Methods

### Strains, plasmids, and media

The *C. neoformans* var. *grubii* strain H99 (serotype A) and derived mutants were used in the experiments. *C. neoformans* deletion mutants lacking Xpp1, Epp1 or both proteins (S1 Table) were from our previous study (5). Strains were maintained on YPD rich medium (1% yeast extract, 2% peptone, 2% dextrose, 2% agar). The hygromycin resistance cassette originated from pJAF15. YPD plates containing hygromycin (200 μg/mL) were used to select transformants. Unless specified otherwise, all chemicals were obtained from Sigma-Aldrich. Iron-chelated dH_2_O was prepared by passage through a column containing Chelex-100 resin (BIORAD Chelex-100) and used in the preparation of low-iron media (LIM) (0.5% glucose, 38 mM L-asparagine, 2.3 mM K_2_HPO_4_, 1.7 mM CaCl_2_•H_2_O, 0.3 mM MgSO_4_•7H_2_O, 20 mM HEPES, 22 mM NaHCO_3_, 1 mL of 1000X salt solution (0.005 g/L CuSO_4_•5H_2_O, 2 g/L ZnSO_4_•7H_2_O, 0.01 g/L MnCl_3_•4H_2_O, 0.46 g/L sodium molybdate, 0.057 g/L boric acid in iron-chelated dH_2_O adjusted to pH 7.4 with 0.4 mg/L sterile thiamine added post-filtering) as described previously (62). For growth assays on fermentative and non-fermentative carbon sources, cells cultured in YPD were transferred to liquid YNB media lacking amino acid supplements and containing 2% glucose, 1% acetate or 1% inositol.

### Construction of deletion mutants

The *vtc4*Δ*epp1*Δ*xpp1*Δ (ΔΔΔ) triple deletion mutant was constructed by homologous recombination using gene-specific cassettes amplified via three-step overlapping PCR method utilizing the primers detailed in S2 Table (63). The resistance marker for hygromycin (HYG) was amplified by PCR using primers ΔΔΔ-2 and ΔΔΔ-5 and the plasmid pJAF15, respectively, as the template. Primers ΔΔΔ-1 and ΔΔΔ-3, as well as ΔΔΔ-4 and ΔΔΔ-6, were employed to amplify the surrounding sequences of the *vtc4* gene, while primers ΔΔΔ-1 and ΔΔΔ-6 were utilized to amplify the deletion construct of the gene, incorporating the resistance marker. The construct was introduced into the *epp1*Δ*xpp1*Δ-4 strain, by biolistic transformation, as described previously (64). Colony PCR was performed to identify positive transformants using primers ΔΔΔ-7N, ΔΔΔ-8N, HYGL and HYGR. Primers used in mutant screening and verification are summarized in S2 Table. Two independent mutants were used for all experiments unless stated otherwise.

### Murine infection and assessment of virulence

Female BALB/c mice, 4-6 weeks old, were obtained from Charles Rivers Laboratories (Pointe-Claire, Quebec, Canada) and housed in the Modified Barrier Facility at the University of British Columbia (UBC). Mice were housed in sterilized cages with microisolator cage tops, maintaining specific-pathogen-free conditions. Clean food and water were provided *ad libitum*. For infections, fungal cells were cultured in 5 mL YPD at 30°C overnight, washed three times with physiological saline, and resuspended in physiological saline (pH 7.4). A cell suspension of 2 x 10^5^ cells in 50 μL was intranasally instilled. The status of the mice was monitored once per day post-inoculating. Mice reaching the experimental endpoint of 15% weight loss were euthanized by an overdose of isoflurane followed by carbon dioxide asphyxiation. Differences in virulence were statistically assessed using log rank tests. For determination of fungal burden, lungs, livers, spleens, kidneys, and livers were excised, weighed, and homogenized in 2 volumes of 1X phosphate-buffered saline using a MixerMill (Retsch, Cole-Parmer, Montreal, Canada). Serial dilutions of the homogenates were plated on YPD plates containing 50 μg/mL chloramphenicol, and colony-forming units were counted after 48 hours at 30°C.

### Ethics statement

This study was carried out in strict accordance with the guidelines of the Canadian Council on Animal Care. The protocol for the virulence assays employing mice was approved by the University of British Columbia Committee on Animal Care.

### Cytometric Bead Array (CBA) Assay

Lungs were excised and collected in PBS containing 2x cOmplete, EDTA-free protease inhibitor cocktail (Roche). Lungs were homogenized using a mixer mill. Supernatants of homogenates were collected and cytokines in the samples were analyzed using BD cytometric Th1/Th2/Th17 kit (BD Biosciences). The kit was used for the simultaneous detected of mouse interleukin-2 (IL-2), interleukin-4 (IL-4), interleukin-6 (IL-6), interferon-γ (IFN-γ), tumor necrosis factor (TNF), interleukin-17A (IL-17A), and interleukin-10 (IL-10) in a single sample. The operations were formed according to the manufacturer’s instructions. Beads coated with seven specific capture antibodies were fixed. Subsequently, 50 μL of the mixed captured beads, 50 μL of the unknown lung supernatant or standard dilution, and 50 μL of phycoerythrin (PE) detection reagent were added to each assay tube and incubated for 2 hours at room temperature in the dark. The samples were washed once with 5 mL of wash buffer and centrifuged. The bead pellet was resuspended in 300 μL buffer. Samples were measured on the CytoFLEX S (Beckman Coulter) Flow Cytometer equipped with four laser lines (405 nm, 488 nm, 561 nm and 633 nm) fitted with filters FITC (525/40), PE (585/42), ECD (600/20). Individual cytokine concentrations were indicated by their fluorescent intensities. Cytokine standards were serially diluted to facilitate the construction of calibration curves, necessary for determining the protein concentrations of test samples.

### Histology

Lungs were harvested and fixed overnight in 10% buffered formalin. Samples were dehydrated, paraffin embedded, sectioned, and stained with either hematoxylin and eosin (H&E), Mayer’s mucicarmine or Movat’s Pentachrome. Images were acquired using a Nikon ECLIPSE Ti2 inverted microscope equipped with a Nikon DS-Ri2 high resolution full-frame sensor microscope camera (Nikon Instruments Inc.). For visualization and image processing, NIS-Elements Viewer software was used.

### Bronchoalveolar lavage (BAL)

Mice were euthanized through an isoflurane overdose, followed by carbon dioxide asphyxiation. Alveolar cells were collected via BAL using catheterization of the trachea and washed three times, with each wash consisting of 1 mL 1X PBS and 2 mM EDTA. The cells were then pelleted, treated with RBC lysis buffer (10 mM Tris, 0.84% NH_4_Cl), and prepared for flow cytometry.

### Tissue processing and analysis of immune cells by flow cytometry

Lungs were perfused with 15-20 mL of ice cold 1 X PBS (pH 7.4) until tissue turned white. Perfused lungs were excised, minced and enzymatically digested in 5 mL of 0.7 mg/mL collagenase IV (Worthington), 50 μg/mL DNAse I (Worthington) in RPMI 1640 for 1 hour at 37°C. Dissociated cells and tissue were passed through a 70 μm strainer, red blood cells were lysed with RBC lysis buffer (10 mM Tris, 0.84% NH_4_Cl) for 5 min at room temperature, and the remaining homogenate was passed through a 35 μm strainer to generate a single cell suspension. Isolated single cells were then incubated with 2.4G2 tissue culture supernatant to block Fc receptors and immunolabeled for cell surface antigens for 20 minutes at 4°C in flow cytometry buffer (PBS, 2% bovine serum albumin, 2 mmol/L EDTA). Multiparameter assessment was performed using Attune NxT Flow cytometer (Thermo Fisher Scientific) and data were analyzed with FlowJo software (TreeStar). The following antibodies against mouse antigens were used for flow cytometry: CD45 (30-F11), CD24 (M1/69), CD11c (N418), CD11b (M1/70), CD64 (X54-5/7.1), Ly6C (HK1.4), Ly6G (1A8), CD4 (RM4-5) from BioLegend; NK1.1 (PK136), SiglecF (1RNM44N), MHCII (M5/114.15.2) from eBioscience; CD3 (145-2C11), CD8 (53-6.7), CD19 (1D3) from AbLab (UBC). Zombie Aqua fixable viability kit was used to identify live cells (BioLegend). Cell numbers were calculated using counting beads (123 count eBeadsTM, eBioscience/Thermo Fisher Scientific). Data were analyzed in GraphPad Prism 10.0, and a two tailed unpaired Student *t* test was used when comparing different biological samples.

### Stress and drug response assays

Exponentially growing cultures of WT or mutant strains were washed, diluted to an initial concentration of 1 x 10^8^ cells/mL in H_2_O, diluted 10-fold serially and 5 μL of each dilution was spotted onto YPD or YNB plates supplemented with the following compounds: 2.5 mM ZnCl_2_, 3.0 mM NiSO_4_, 1.875 mM MnSO_4_, 0.5 M CaCl_2_, 1 mg/mL Caffeine, 1 mg/mL CFW, 1% Congo Red, 0.5% SDS, 1 mM or 3 mM H_2_O_2_, 15 mM Salicylhydroxamic acid (SHAM), 10 mM or 20 mM KCN, 5 μg/mL Antimycin A, 5 μM Myxothiazol, 50 μg/mL Rotenone, 0.25 mM Paraquat dichloride, 25 μM Plumbagin (Cedarlane Labs), or 1 mM tert-Butyl hydroperoxide (tBuOOH). Plates were incubated for 2-5 days at 30°C, 37°C or 39°C and photographed.

### Macrophage uptake and survival assays

J774A.1 murine macrophage-like cells (ATCC, Manassas, Virginia) were routinely cultured in high glucose Dulbecco’s Modified Eagle Medium (Fisher Scientific) supplemented with 10% heat-inactivated fetal bovine serum (FBS), 2 mM L-glutamine (Gibco) and utilized within 20 passages after thawing from lN_2_. Bone marrow-derived macrophages (BMDMs) were generated by culturing bone marrow cells with L-929 cell conditioned medium (LCCM) for 6 days. The cultivation of macrophages was performed in DMEM supplemented with LCCM, 20% bovine serum albumin, 1 mM Na-pyruvate and 2 mM L-glutamine at 37°C, 5% CO_2_. Phagocytosis and intracellular replication of WT and mutant strains in the J774A.1 mouse macrophage-like cell line and BMDMs were determined as previously described (65). Briefly, macrophage cells in 24-well plates were stimulated with 150 ng/mL phorbol myristate acetate (PMA) for 1 hour prior to infection. Cells of the wild type or mutant strains were opsonized for 1 hour with the monoclonal antibody 18B7. Stimulated macrophages were infected with 2 x 10^6^ opsonized yeast cells (10:1 multiplicity of infection; yeast: macrophage) for 2 and 24 hours at 37 °C, 5% CO_2_. Non-internalized yeast cells were washed off three times with 1 X PBS. To determine the number of internalized yeast cells, macrophages were lysed with sterile distilled H_2_O and lysates were plated on YPD agar for CFU counts at both 2 and 24 hours.

BMDM and J774A.1 macrophage cells were seeded in eight-well chamber slides in DMEM medium and allowed to adhere firmly overnight. The next day, macrophage cells were washed and maintained in serum-free DMEM for infection with opsonized yeast cells. For the analysis of macrophage uptake, live differential interference contrast (DIC) images were taken 2 hours post-infection and at least 300 macrophages were counted per strain for reach experiment. For the analysis of intracellular survival, non-internalized yeast cells were washed off three times with 1X PBS and live DIC images were taken 24 hours post-infection.

### Fluorescence microscopy

For chitin detection, cells were grown overnight in YPD, washed twice in 1 X PBS, diluted to an OD_600_nm of 1 and stained with calcofluor white (CFW) in PBS for 15 minutes at room temperature in the dark. For visualization of mitochondria and analysis of mitochondrial morphology, *C. neoformans* cells were grown overnight in YPD, transferred to YNB with or without 1 mM H_2_O_2_ for 1 hour. After treatment with H_2_O_2_, cells were washed with 1X PBS before staining with mitochondria specific dyes nonyl acridine orange (NAO, 200 ng/μL) or MitoTracker Orange CMTMRos (200 nM) for 30 minutes in dark at 30°C. Cells were collected, washed with 1X PBS three times and kept on ice before microscopy. Differential interference contrast (DIC) and fluorescence imaging were performed with a wide-field fluorescence microscope (Zeiss Axiovert 200) coupled with a CMOS camera (ORCA-Flash4.0 LT; Hamamatsu Photonics). For polyP granule detection, *C. neoformans* cells were grown overnight in YPD, washed three times in 1 X PBS, diluted to an OD600nm of 1 and stained with 4′,6-diamidino-2-phenylindole (DAPI) at a concentration of 100 μg/mL for 30 minutes at room temperature. After staining, the cells were washed two times with 1 X PBS. Fluorescence microscopy images were acquired using the wide-field fluorescence equipment described above using the BrightLine® full multiband filter set (AVR Optics).

### Flow cytometry analysis measurements for intracellular ROS accumulation

Flow cytometric measurements were performed using CytoFLEX S (Beckman Coulter) Flow Cytometer equipped with four laser lines (405 nm, 488 nm, 561 nm and 633 nm) fitted with filter FITC (525/40). The number of cells measured per experiment was set to 30,000-40,000 unless otherwise stated. For the ROS sensitivity analysis, cells were grown overnight at ∼200 rpm and 30°C in YPD media and washed twice with 1X PBS. Subsequently, 0.4 OD cells were grown in 25 mL of YNB media for 3-4 hours at 200 rpm and 30°C. Cells were collected and then treated in YNB with or without hydrogen peroxide (H_2_O_2_, 5 mM) for 1 hour. After ROS treatment, cells were collected and stained with 2ʹ,7ʹ-Dichlorofluorescein Diacetate (DCFDA, 16 μM) for 1 hour at 30°C before cytometric analysis.

### Assessment of polyP accumulation

The amount of intracellular polyP was determined as previously described (5,34). Briefly, cells were grown in YPD overnight at 30°C. RNA was extracted using a citrate buffer and bead beating in a bead mill to rupture the cells and release RNA and polyphosphate. Subsequently, 10 μg of total RNA was loaded onto a native DNA polyacrylamide gel and subjected to electrophoresis in 1× Tris-borate-EDTA (TBE) buffer. The RNA and polyphosphate were then fixed with acetate, stained with toluidine blue O, and destained in acetate, following previously established methods. 10 μg of polyphosphate type 700 (P700; Kerafast) was used as molecular marker control.

### Assessment of virulence factor production

Elaboration of polysaccharide capsule was examined by differential interference microscopy (DIC) using Zeiss Plan-Apochromat 100x/1.46 oil lens on a Zeiss Axioplan 2 microscope coupled with a CMOS camera (ORCA-Flash4.0 LT; Hamamatsu Photonics) after incubation for 48 hours at 30 °C in defined low-iron medium (LIM). Cells were stained with India ink to visualize polysaccharide capsule. Melanin production was assessed through growth on L-3,4- dihydroxyphenylalanine (L-DOPA) plates or liquid medium containing 0.1% glucose, as previously described (66).

### Sample Preparation and Scanning Electron Microscopy

*C. neoformans* cells grown in low iron, capsule inducing media (CIM) for 48 hours at 30°C were washed twice in 1X PBS (pH 7.4) and fixed for 1 hour in 2.5% glutaraldehyde in 0.1 M sodium cacodylate buffer (pH 7.4) at room temperature. Subsequently, the fixed cells were washed twice in 1 X PBS. 20 μL of fixed cell solution was deposited on poly-L-lysine coated coverslips (BioCoat Ref#354085). Samples were gradually dehydrated in ethanol series: 50, 60, 70, 80, 90 and three subsequent 100%. Dehydrated samples were dried using a Tousimis Samdri-795 critical point dryer before coating with 10 nm iridium using a Leica EM MED020 sputter coater. Images were acquired using a Helios NanoLab 650 dual beam SEM (Thermofisher, MA) with an Everhart-Thornley detector at a voltage between 10-15 kV and working distance of 4 mm.

### Statistical Analysis

Statistical analysis was performed using unpaired student *t* tests, Kruskal-Wallis, Mann-Whitney, or one-way ANOVA. All statistical tests were conducted using GraphPad Prism software.

## Acknowledgements

We recognize the contributions of Andy Johnson from the UBC Flow Cytometry Core and Marcos Aguiar, Angel Chu, Miaoran Li, and animal care technicians at the UBC Modified Barrier Facility. We acknowledge the UBC Centre for High-Throughput Analysis for their support in electron microscopy imaging. We also thank Nigel O’Neil, Manisha Dosanjh, Tenanye Haglund, Melissa Lagace and Haohua Li for technical assistance.

## Supporting information

**S1 Fig.** Infiltration of immune cells in the lung and BAL of mice infected with *C. neoformans*. Immune cell analysis in the BAL fluid or lung tissue of BALB/c mice infected with WT, *vtc4*Δ, or *xpp1*Δ*epp1*Δ, or treated with physiological saline at 7 days post infection. Gating strategy was described previously in Figure 4A. Total number of adaptive B cells (**A**), Ly6C^-^macrophages, including alveolar macrophages (AM) and interstitial macrophages (IM) (**B**), and innate immune cells, including cDC1, cDC2, NK and NKT cells (**C**) in the lung and BAL. Data are presented as mean ± SD and representative of at least 2 independent experiments for each time point (n = 4 mice/time point). Significance indicated as *, *P* < 0.05; **, *P* < 0.01; ***, *P* < 0.001; unpaired Student *t* test.

**S2 Fig.** The *vtc4*Δ*epp1*Δ*xpp1*Δ triple deletion mutant does not have detectable polyP and exhibits sensitivity to zinc. **(A)** Representative widefield fluorescence micrographs showing polyP granules in WT, *xpp1*Δ, *epp1*Δ, *xpp1*Δ*epp1*Δ, or *vtc4*Δ*epp1*Δ*xpp1*Δ (ΔΔΔ) strains. Indicated strains were stained with 4′,6- diamidino-2-phenylindole (DAPI, 100 μg/mL) for 30 minutes. Scale bar 5 μm. **(B)** Representative native polyacrylamide gel stained with toluidine blue O showing polyP accumulation in indicated strains. A marker of polyP type 700 (10 μg) was loaded alongside 10 μg of RNA extracted from each strain (cultured overnight in YPD). Data representative of at least three independent experiments. **(C)** Indicated strains were serially diluted and spotted onto solid YPD agar with or without 2.5 mM ZnCl_2_, 3.0 mM NiSO_4_, 1.875 mM MnSO_4_, or 0.5 M CaCl_2_. The plates were then incubated at 30 °C for 2-5 days before being photographed.

**S3 Fig.** Mutants deficient in polyP synthesis and mobilization are not sensitive to cell wall stress. Indicated strains were serially diluted and spotted onto solid YPD agar with or without 1 mg/mL Caffeine, 1 mg/mL CFW or 1% Congo Red. The plates were then incubated at 30 °C for 5 days before being photographed.

**S4 Fig.** Loss of polyP synthesis and mobilization does not confer sensitivity to all ROS or ETC stress agents. Indicated strains were serially diluted and spotted onto solid YPD agar with or without 0.25 mM Paraquat dichloride, 25 μM Plumbagin, 1 mM tert-Butyl hydroperoxide (tBuOOH) **(A),** 5 μg/mL Antimycin A, 5 μM Myxothiazol, or 50 μg/mL Rotenone **(B)**. The plates were then incubated at 30°C or 37°C for 2-5 days before being photographed. **(C)** Liquid growth assays on YNB medium supplemented with 2% glucose, or 1% acetate for the indicated strains. *C. neoformans* yeast cells were grown in YNB without amino acids with 2% glucose or 1% acetate in test tubes containing 5 mL of the indicated growth medium. The tubes were incubated at 30°C, 200 rpm for 120 hours and optical densities (OD600) were measured every 24 hours. Data are presented as mean ± SEM representative of at least 3 independent experiments.

